# Constitutively enhanced genome integrity maintenance and direct stress mitigation characterize transcriptome of extreme stress-adapted *Arabidopsis halleri*

**DOI:** 10.1101/859249

**Authors:** Gwonjin Lee, Hassan Ahmadi, Julia Quintana, Lara Syllwasschy, Nadežda Janina, Veronica Preite, Justin E. Anderson, Björn Pietzenuk, Ute Krämer

## Abstract

Heavy metal-rich toxic soils and ordinary soils are both natural habitats of *Arabidopsis halleri*. The molecular divergence underlying survival in sharply contrasting environments is unknown. Here we comparatively address metal physiology and transcriptomes of *A. halleri* originating from the most highly heavy metal-contaminated soil in Europe, Ponte Nossa (Noss/IT), and from non-metalliferous (NM) soil. Noss exhibits enhanced hypertolerance and attenuated accumulation of cadmium (Cd), and transcriptomic Cd responsiveness is decreased, compared to plants of NM soil origin. Among the condition-independent transcriptome characteristics of Noss, the most highly overrepresented functional class of “meiotic cell cycle” comprises 21 transcripts with elevated abundance in vegetative tissues, in particular *Argonaute 9* (*AGO9*) and the synaptonemal complex transverse filament protein-encoding *ZYP1a/b*. Increased *AGO9* transcript levels in Noss are accompanied by decreased long terminal repeat retrotransposon expression, and are shared by plants from milder metalliferous sites in Poland and Germany. Expression of *Iron-regulated Transporter* (*IRT1*) is very low and of *Heavy Metal ATPase 2* (*HMA2*) strongly elevated in Noss, which can account for its specific Cd handling. In plants adapted to the most extreme abiotic stress, broadly enhanced functions comprise genes with likely roles in somatic genome integrity maintenance, accompanied by few alterations in stress-specific functional networks.

## Introduction

Heavy metals, such as zinc (Zn), cadmium (Cd) and lead (Pb), are naturally omnipresent in the biosphere at very low levels. Their concentrations can be regionally elevated and are extremely high locally at rare sites as a result of geological anomalies. Human activities, for example ore mining, metallurgical processing and sewage sludge deposition, contribute to anthropogenic contamination of the biosphere with heavy metals, threatening environmental and human health (Nriagu and Pacyna, 1988, Lamas *et al*., 2016). Heavy metal-enriched habitats often host characteristic ecological communities, especially where human activities have introduced extremely high, toxic levels of heavy metals only recently after the beginning of the industrial revolution (Ernst, 1974). Here we use comparative transcriptomics to uncover the molecular basis of local adaptation to an extremely highly heavy metal-contaminated soil in a land plant as a complex multi-cellular eukaryotic organism.

The Brassicaceae species *Arabidopsis halleri*, in the sister clade of the genetic model plant *Arabidopsis thaliana*, is an emerging perennial model plant (Krämer, 2015, Honjo and Kudoh, 2019, Nagano *et al*., 2019). *A. halleri*, a perennial stoloniferous obligate outcrosser, has repeatedly colonized highly heavy metal-contaminated, so-called metalliferous (M) soils (Ernst, 1974, Bert *et al*., 2000, Stein *et al*., 2017). Natural populations of *A. halleri* are also found on non-contaminated (non-metalliferous, NM) soils. Irrespective of its habitat soil type, *A. halleri* is a Zn and Cd hyperaccumulator (Ernst, 1974, Bert *et al*., 2000, Stein *et al*., 2017). The about 750 known hyperaccumulator plant species are land plants of which at least one individual was identified to accumulate a metal or metalloid in its above-ground tissues to concentrations more than one order of magnitude above critical toxicity thresholds of ordinary plants, at its natural site of growth in the field (Reeves *et al*., 2018). Hyperaccumulation in plants can act as an elemental defense against biotic stress (Kazemi-Dinan *et al*., 2014), and the associated species-wide heavy metal tolerance can apparently facilitate the colonization of M soils by *A. halleri* (Pauwels *et al*., 2005, Meyer *et al*., 2010).

Cross-species comparative transcriptomics studies of *A. halleri* and *A. thaliana* established tens of candidate metal homeostasis genes for roles in metal hyperaccumulation or metal hypertolerance, which encode various transmembrane transporters of metal cations and isoforms of a metal chelator biosynthetic enzyme (Becher *et al*., 2004, Weber *et al*., 2004, Talke *et al*., 2006). Although the generation of transgenic plants remains technically demanding and time-consuming in *A. halleri*, subsequent work demonstrated functions in metal hyperaccumulation or hypertolerance for some of these genes. The key locus making the largest known contribution to both metal hyperaccumulation and hypertolerance in *A. halleri* is *Heavy Metal ATPase 4* (*HMA4*), which encodes a plasma membrane P_1B_-type metal ATPase mediating cellular export for xylem loading of Zn and Cd in the root (Talke *et al*., 2006, Hanikenne *et al*., 2008). Transcript levels of *HMA4* are substantially higher in *A. halleri* than in *A. thaliana* independently of cultivation conditions, as was also observed for tens of other candidate genes. A combination of modified *cis*-regulation and gene copy number expansion accounts for high expression of *HMA4* (Hanikenne *et al*., 2008). Examples of other candidate genes with experimentally demonstrated functions in *A. halleri* include *Nicotianamine Synthase 2* (*NAS2*) (Deinlein *et al*., 2012) contributing to Zn hyperaccumulation, *Metal Transport Protein 1* (*MTP1*) acting in Zn tolerance (Dräger *et al*., 2004) and *Ca*^*2+*^*/H*^*+*^ *exchanger 1* (*CAX1*) functioning to enhance Cd tolerance (Baliardini *et al*., 2015). While metal hyperaccumulation and hypertolerance are species-wide traits in *A. halleri*, there is additionally a large extent of within-species variation in the levels of metal accumulation, metal tolerance and gene expression (Meyer *et al*., 2015, Stein *et al*., 2017, Corso *et al*., 2018, Schvartzman *et al*., 2018).

Within-species transcriptomic differences between *A. halleri* populations originating from M and NM soils can provide insights into the molecular and physiological alterations associated with the natural colonization of M soils, but they have hardly been explored thus far. In order to address adaptation to extremely high heavy metal levels in a metalliferous soil in *A. halleri*, we focused here on the population at the most highly heavy metal-contaminated *A. halleri* site in Europe at Ponte Nossa (Noss/IT) according to a large-scale field survey of 165 European populations (Stein *et al*., 2017). Noss individuals were more Cd-tolerant and accumulated lower levels of Cd than both closely related *A. halleri* from a population on NM soil in the vicinity, Paisco Loveno (Pais), and more distantly related *A. halleri* from Wallenfels (Wall) on NM soil in Germany. Further contrasting with the less Cd-tolerant plants of NM soil origin, transcriptomic Cd responses including the activation of Fe deficiency responses were attenuated in the highly Cd-tolerant population. Instead, a large number of transcripts differed in abundance in Noss (M soil origin) from both Pais and Wall plants (NM soil origins) irrespective of cultivation conditions. Overrepresentation analysis indicated that differences comprised primarily the global activation of meiosis- and genome integrity maintenance-related functions in somatic tissues of Noss, accompanied by pronouncedly altered levels of few metal homeostasis transcripts known for central roles in Cd accumulation and basal Cd tolerance of *A. thaliana*. Our results suggest the possibility of roles in somatic tissues of *A. halleri* (Noss) for some genes functions known to operate in meiosis of *A. thaliana*. This work provides insights into how land plants cope with rapid anthropogenic environmental change and extreme abiotic stress levels.

## Materials and Methods

For detailed Materials and Methods see Supplemental Information.

### Plant material and growth conditions

Cuttings were made of *A. halleri* ssp. *halleri* O’Kane & Al-Shehbaz individuals originating from the field (Stein *et al*., 2017), followed by hydroponic pre-cultivation for 4 weeks to obtain rooted vegetative clones. Experiments were conducted in 1x modified Hoagland’s solution (Becher *et al*., 2004) with weekly exchange of solutions. In Cd tolerance assays, plants were sequentially exposed to step-wise increasing concentrations of CdSO_4_ (0, 25, 50, 75, 100, 125, 150, 175, 200, 250, 300, 350, 400 and 450 μM) once per week in a growth chamber (10-h light at 90 μmol photons m^-2^ s^-1^, 22°C/ 14-h dark at 18°C, 65% constant relative humidity; *n* = 6 to 9 per population, with one to three vegetative clones for each of three to six genotypes per population). Cd tolerance index was quantified as EC_100_, the effective Cd concentration causing 100% root growth inhibition (Schat and Ten Bookum, 1992). For transcriptome sequencing and multi-element analysis (Table S4), 1x modified Hoagland’s solution was supplemented with either 2 or 0 (control) μM CdSO_4_, with cultivation in 16-h light (90 μmol photons m^-2^ s^-1^, 22°C)/ 8-h dark (18°C) cycles at 60% constant relative humidity for 16 d.

### Multi-element analysis and transcriptome sequencing

For multi-element analysis, apoplastically bound metal cations were desorbed from freshly harvested roots (Cailliatte *et al*., 2010)(see Supplemental Information). All tissues were washed twice in ddH_2_O before drying, digestion and multi-element analysis as described (Stein *et al*., 2017). For transcriptome sequencing, root and shoot tissues were harvested separately 7 h after the onset of the light period, frozen immediately in liquid nitrogen, pooled from six replicate clones per genotype, treatment and experiment, and stored at -80°C. Total RNA was extracted from aliquots of tissue homogenates using the RNeasy Plant Mini Kit (Qiagen, Hilden, Germany). One μg of total RNA was used in NEBNext Ultra II RNA Library Prep Kit for Illumina (New England Biolabs, Frankfurt, Germany), followed by sequencing to obtain 14 to 29 mio. 150-bp read pairs per sample that passed quality filters (Novogene, Hongkong).

### Sequence data analysis

Reads were mapped to the *A. halleri* ssp. *gemmifera* (Matsumura) O’Kane & Al-Shehbaz accession Tada mine (W302) reference genome (Briskine *et al*., 2017) using HISAT2 version 2.1.0 (Kim *et al*., 2015). This reference genome contains a more complete set of *A. halleri* coding regions than the *A. lyrata* reference genome, and its use resulted in considerably higher mapping rates. After multiple mapping error correction according to COMEX 2.1 (Pietzenuk *et al*., 2016), the number of fragments per gene were determined using Qualimap2 (Okonechnikov *et al*., 2016), followed by principal component analysis (PCA) and differential gene expression analysis using the R package *DESeq2* (Love *et al*., 2014). Clustering was performed using the R package *pheatmap*. For gene ontology (GO) term enrichment analyses *A. halleri* gene IDs were converted into *A. thaliana* TAIR10 AGI codes, which were then used in hypergeometric distribution estimation through the function g:GOSt built in g:Profiler (Reimand *et al*., 2016). Transposable element (TE) transcript levels were quantified based on Reads Per Kilobase per Million reads (Mortazavi *et al*., 2008) after mapping to the *Arabidopsis lyrata* MN47 reference genome with TE annotations (Pietzenuk *et al*., 2016). This was necessary because the *A. halleri* ssp. *gemmifera* reference genome lacks contiguity outside coding sequences and in repetitive regions and its TE annotations are less thoroughly curated. Reads were used to reconstruct cDNA variants with Integrated Genome Viewer (Robinson *et al*., 2017). Multiple comparisons of means were conducted using the *stats* and *agricolae*, with normality and homoscedasticity tests in the *car*, packages in R (R_Core_Team, 2013).

### Validation by RT-qPCR and immunoblots

We used aliquots of homogenized tissues as frozen during harvest. Real-time RT-qPCR reactions were run in a 384-well LightCycler®480 II System (Roche Diagnostics, Mannheim, Germany) using the GoTaq qPCR Mastermix (Promega, Walldorf, Germany) and cDNAs synthesized from DNase-treated total RNA using the RevertAid First Strand cDNA Synthesis Kit (Thermo Fisher Scientific, Schwerte, Germany). Immunoblots were carried out following SDS-PAGE (Lämmli, 1970) using anti-*At*IRT1 (AS11 1780, Lot number 1203; Agrisera, Vännas, Sweden), HRP-conjugated secondary antibodies (Thermo Fisher Scientific) and detection using the ECL Select Western Blotting Detection Reagent (GE Healthcare, Little Chalfont, England) and a Fusion Fx7 GelDoc (Vilber Lourmat, Eberhardzell, Germany).

## Results

### Choice of populations and comparative physiological characterization

In a field survey of 165 *Arabidopsis halleri* sites across Europe, average soil Cd concentrations were by far the highest at Ponte Nossa/IT (Noss) in both the total and hydrochloric acid-extractable soil fractions (900 and 203 mg Cd kg-1 soil, respectively; second highest were soils of far lower concentrations of 130 and 87 mg Cd kg^-1^ soil, respectively)(Stein *et al*., 2017). Therefore, we chose the *A. halleri* population at this M site as the focal population in this comparative study, in combination with two populations at NM sites for comparison, one from the geographically closest site near Paisco Loveno (Pais/IT) approximately 38 km northeast of Noss, and one near Wallenfels (Wall/DE) about 520 km to the North. Average soil total Cd concentrations in close proximity of the *A. halleri* individuals studied here were 700-fold higher at Noss than at Pais (Table S1). Earlier comparative transcriptomics studies of Zn/Cd hyperaccumulator plant species included individuals originating from geographically distant M sites only, for example in Italy and Poland for *A. halleri*, and they were not designed to compare between M and NM sites (Corso *et al*., 2018, Schvartzman *et al*., 2018, Halimaa *et al*., 2019).

In order to test for adaptation of the Noss population to locally extremely high soil Cd levels, we quantified Cd tolerance of individuals collected at each of the three sites in hydroponic culture system in a growth chamber. Based on the proportion of individuals maintaining root growth in a sequential exposure test with stepwise increasing Cd concentrations, individuals from Noss were the most tolerant of the three tested populations (Fig. 1a, Table S2, Fig. S1). Following a different approach for quantifying metal tolerance (Schat and Ten Bookum, 1992), mean EC_100_, the effective Cd concentration causing 100% root growth inhibition in a given individual, was significantly higher for Noss (344 ± 98 μM Cd) than for Pais (219 ± 66 μM Cd) and Wall (221 ± 56 μM Cd) (Fig. 1b). In line with species-wide Cd hypertolerance of *A. halleri*, 100% root growth inhibition was previously reported to occur approximately at 30 µM Cd in *A. thaliana* (Becher, 2003).

**Figure 1.**
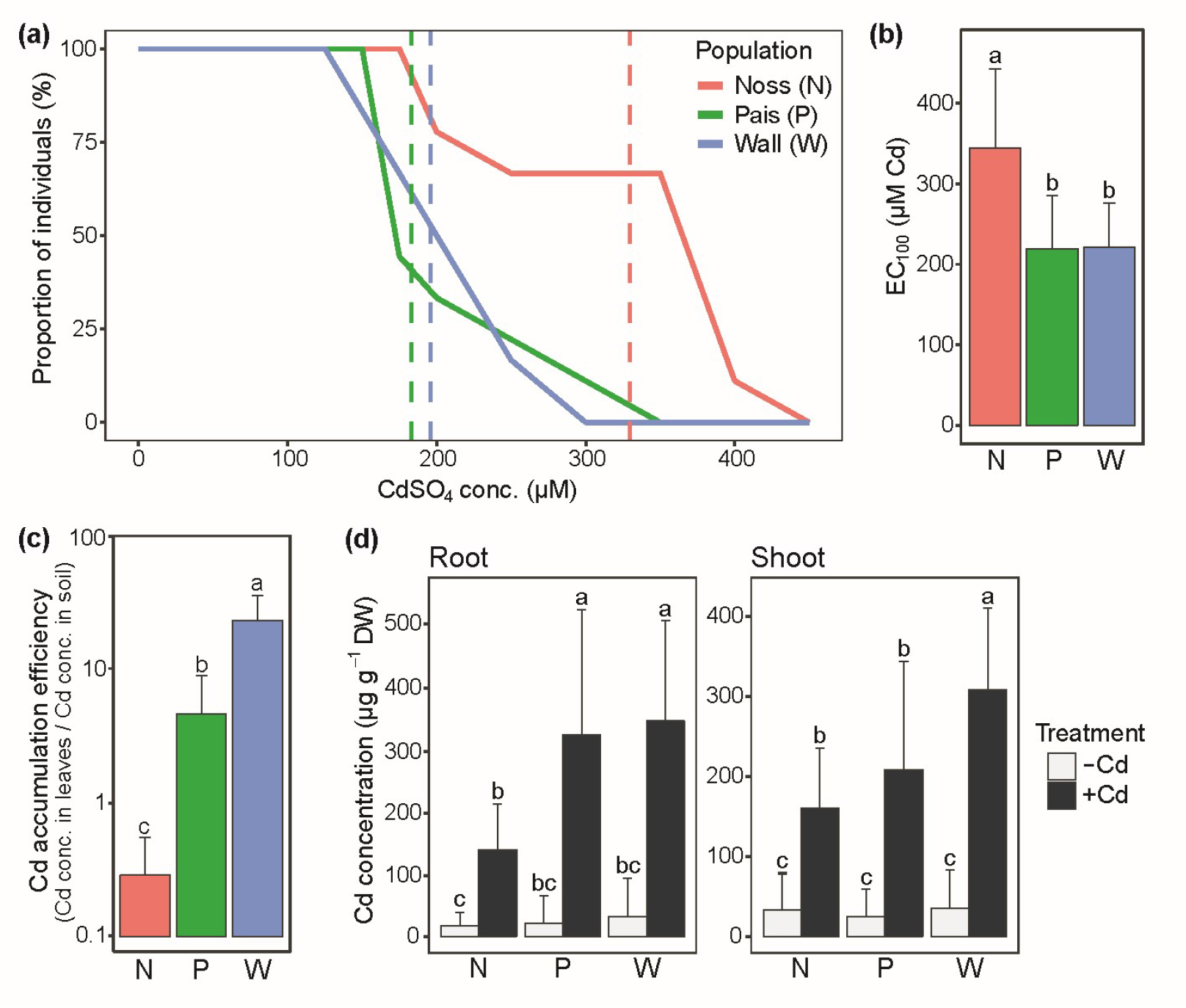
Cadmium tolerance and accumulation in three populations of *A. halleri*. (a) Proportion of individuals maintaining root growth (elongation) as a function of Cd concentration. Dashed vertical lines indicate ED_50_, the effective dose required to reach a proportion of 50% according to a dose-response model (see Table S2, Figure S1). (b) Cd tolerance index. Bars represent mean ± SD of EC_100_ (effective Cd concentration causing 100% root growth inhibition) in a given plant individual (Schat and Ten Bookum, 1992). Plants were sequentially exposed to stepwise increasing concentrations of CdSO_4_ in hydroponic culture every week (*n* = 6 to 9 per population, with one to three vegetative clones of each three to six genotypes per population; (a, b)). (c) Cd accumulation efficiency at the site of origin calculated from published field survey data (bars represent means ± SD of *n* = 11 to 12 individuals in the field; units employed, µg Cd g^-1^ dry biomass in leaves, µg total Cd g^-1^ dry mass in soil; data from Stein et al. 2017). (d) Cd concentration in root and shoot tissues of hydroponically cultivated plants. Shown are mean ± SD (*n* = 12 to 20 clones per population comprising both genotypes, from all three replicate experiments; see Supplemental Table 1). Four-week-old vegetative clones were exposed to 0 (-Cd) and 2 µM CdSO_4_ (+Cd) in hydroponic culture for 16 d alongside the plants cultivated for RNA-seq. Different characters denote statistically significant differences between means based on two-way ANOVA, followed by Tukey’s HSD test (Log-transformed data were used in (c)) (*P* < 0.05; see Supplemental Table 3, for nested ANOVA of genotypes within a population (d)).

In natural populations in the field, Cd concentrations accumulated in leaves of *A. halleri* at Noss were at least 20-fold higher than at Pais and 3-fold higher than at Wall (Table S1)(Stein *et al*., 2017). By contrast, Zn concentrations in leaves of field-collected individuals were similar in all three populations, despite vastly differing soil Zn levels (see Table S1)(Stein *et al*., 2017). The leaf:soil ratio of Cd concentrations (Cd accumulation efficiency) in the field was highest in the Wall population from north of the Alps and lowest in Noss, suggesting attenuated leaf Cd accumulation at Noss compared to both Pais and Wall (Fig. 1c). Upon exposure to a sub-toxic concentration of 2 μM CdSO_4_ in hydroponic culture for 16 d, Cd concentrations in roots of Noss (140 ± 74 μg Cd g^−1^ DW) were only 43% and 40% of those in Pais and Wall, respectively (Fig. 1d, Table S3). Leaf Cd concentrations were also lower in Noss (160 ± 74 μg Cd g^−1^ DW) than in Wall (about 52% of Wall; *P* < 0.05), and at about 76% of those in Pais (n.s.). We additionally observed between-population differences in the accumulation of other nutrients (Fig. S2). Upon Cd exposure, Cu concentrations were higher, and Mn concentrations were lower than under control conditions in roots of Pais and Wall, but not in roots of Noss (Fig. S2a and d). Fe concentrations were higher in roots of Noss and increased further following exposure to Cd, in contrast to the two populations from NM soils (Fig. S2e). In shoots of Noss, Fe concentrations were about twice as high (*ca*. 75 µg Fe g^-1^ DW) than in the shoots of the other two populations under both control and Cd exposure conditions (Fig. S2e). Taken together, these results showed that the Noss population of *A. halleri* is more tolerant to Cd than both Pais and Wall, supporting adaptation to the composition of its local soil. This was accompanied by attenuated Cd accumulation and a distinct profile of tissue concentrations of micronutrient metals in Noss under standardized growth conditions in hydroponic culture.

### Comparative transcriptomics

Transcriptome sequencing was carried out for a total of 72 samples, with three independent repeats of each experiment comprising root and shoot tissues of two genotypes from each of the three populations grown in control and Cd-amended medium for 16 d (0 and 2 μM CdSO_4_; Table S4; note that this Cd concentration is far sub-toxic in all *A. halleri* populations under investigation and was chosen to elicit possible responses without causing any toxicity symptoms). Principal component analysis (PCA) suggested predominant differences between root and shoot transcriptomes (Fig. S3a). In both roots and shoots, transcriptomes of the geographically neighboring populations Noss and Pais grouped closer together in the first principal component by comparison to the geographically distant Wall population (Fig. S3b and c). This suggested that individuals from Noss and Pais are genetically more closely related to one another than either of these to individuals from Wall. Based on genotyping-by-sequencing of more than 800 European *A. halleri* individuals, we established that Noss and Pais are both in the Central Alpine clade of *A. halleri*, whereas Wall is in the distinct Central European clade (Anderson *et al*., manuscript in preparation).

Upon long-term exposure to a low level of Cd as applied here, transcriptomic differences compared to controls untreated with Cd comprised a few hundred genes in both the Pais and the Wall population originating from NM sites (Fig. 2a and b, Figs. S4 and S5, Dataset S1). By contrast, in the more Cd-tolerant population Noss, no single transcript responded in abundance to Cd exposure under the employed conditions in either roots or shoots. This observation is relevant because it argues against the possible hypothesis that adaptation of *A. halleri* to high-Cd soils involves an enhanced sensitivity of transcriptional responsiveness to Cd. In order to identify candidate transcripts that may contribute to the ability of the Noss population to persist on extremely highly heavy metal-contaminated soil, we next examined the genes showing differential expression between populations (Fig. 2c and d, Dataset S2). As an initial set of candidate genes, we focused on those transcripts showing differential abundance between Noss (M) and both of the populations from NM soils, Pais and Wall. This comparison identified differential transcript levels for as many as 1,617 genes (801/599/217, higher in Noss/ lower in Noss / intermediate in Noss) in roots and 1,638 genes (797/604/237) in shoots under control conditions (under Cd exposure 848/697/236 and 1140/707/339 in roots and shoots, respectively) (surrounded by a red line in Fig. 2c and d).

**Figure 2.**
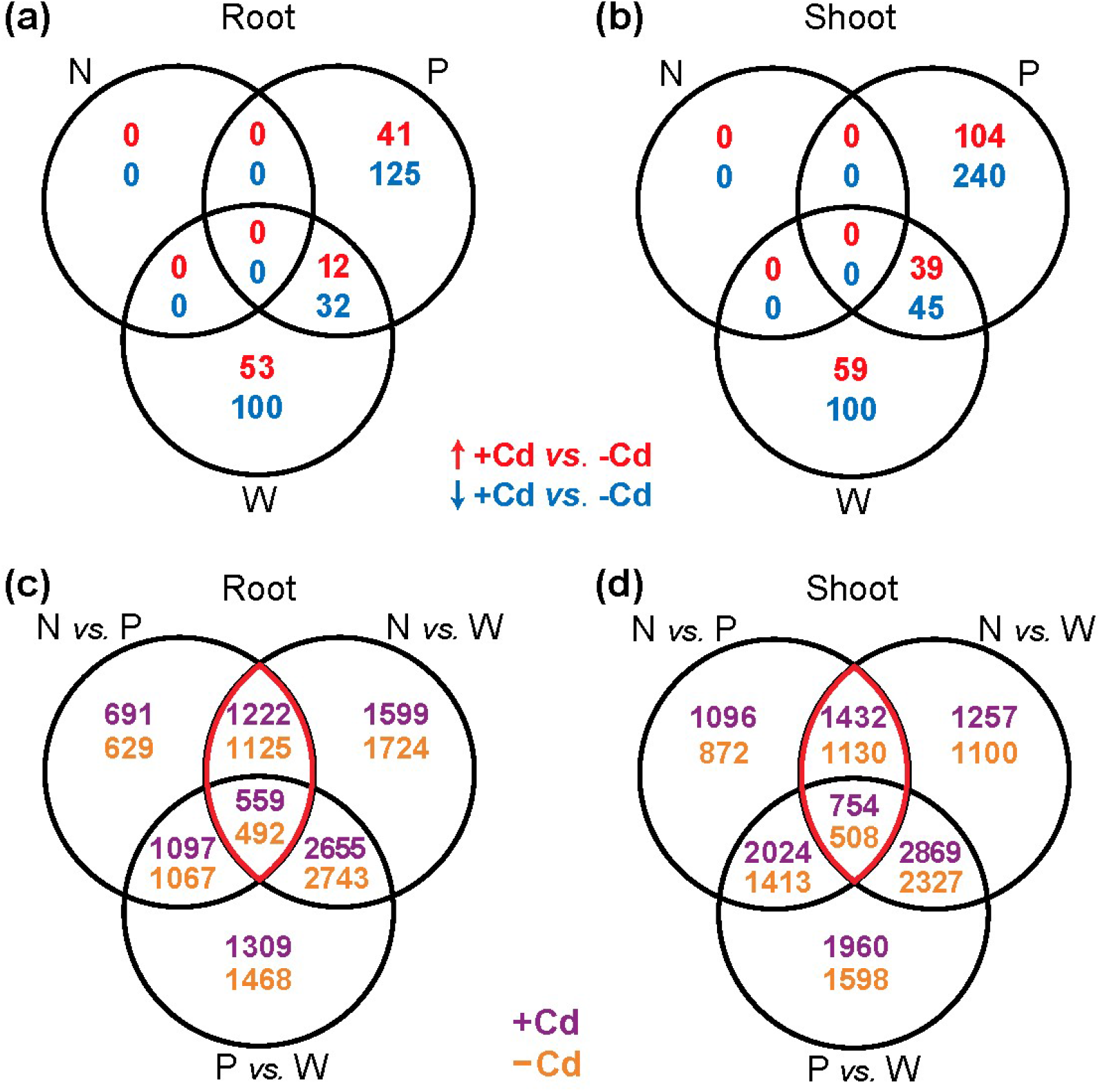
Between-population differences according to transcriptome sequencing. (a, b) Venn diagrams show the numbers of genes exhibiting differential transcript abundances under exposure to Cd (+Cd) compared to control conditions (-Cd) for the three populations Noss (N), Pais (P) and Wall (W) in root (a) and shoot (b) tissues. Upregulation (red): Log_2_(fold change +Cd *vs*. –Cd) = Log_2_FC > 0.5; downregulation (blue): Log_2_FC < -0.5; all with adjusted *P*-value < 0.05. (c, d) Venn diagrams show the numbers of genes exhibiting differential transcript abundance between populations in root (c) and shoot (d) tissues of plants cultivated under Cd exposure (violet) or control conditions (orange) (|Log_2_FC| > 0.5; adjusted *P*-value < 0.05). Red lines surround genes differentially expressed between Noss and both populations of NM soils (Pais and Wall). Four-week-old vegetative clones were exposed to 2 µM Cd or 0 µM Cd (controls) in hydroponic culture for 16 d before harvest. Data are from three independent experiments (repeats), with two genotypes per population and material pooled from six replicate vegetative clones (three hydroponic culture vessels) per genotype in each experiment.

### Candidate genes differentially expressed in Noss

We subjected the genes expressed either at higher or lower levels in Noss compared to both Pais and Wall (DEGs; 0 µM Cd; Dataset S3) to a Gene Ontology Term (GO) enrichment analysis (Dataset S4). Meiotic cell cycle (GO:0051321) was the most significantly enriched in the group of genes that are more highly expressed in both root and shoot of Noss (Fig. 3a and b, Table 1). Among the genes within the GO term meiotic cell cycle, transcript levels of *ARGONAUTE 9* (*AGO9*) were substantially higher in both root and shoot tissues of the Noss in comparison to both the Pais and Wall populations (Fig. 3c and d, Table S5; Log_2_FC = 6.6 and 4.4 to 4.6 in root and shoot, respectively; see Dataset S2). By contrast, present knowledge suggests that the expression of *AGO9* in *A. thaliana* is restricted to only a few cells during the early stages of meiotic megaspore formation (Duran-Figueroa and Vielle-Calzada, 2010, Olmedo-Monfil *et al*., 2010). Amino acid sequence alignment of AGO proteins in *A. halleri* and *A. thaliana* (Fig. S6) confirmed that *Ah*AGO9 is the closest *A. halleri* homologue of *At*AGO9 in clade 3 of ARGONAUTE proteins (AGO4, -6, -8 and -9)(Vaucheret, 2008). Sequence read coverage for *AGO9*, obtained from gDNA of Noss and Pais, suggested that *AGO9* is a single-copy gene (Table S6).

**Figure 3.**
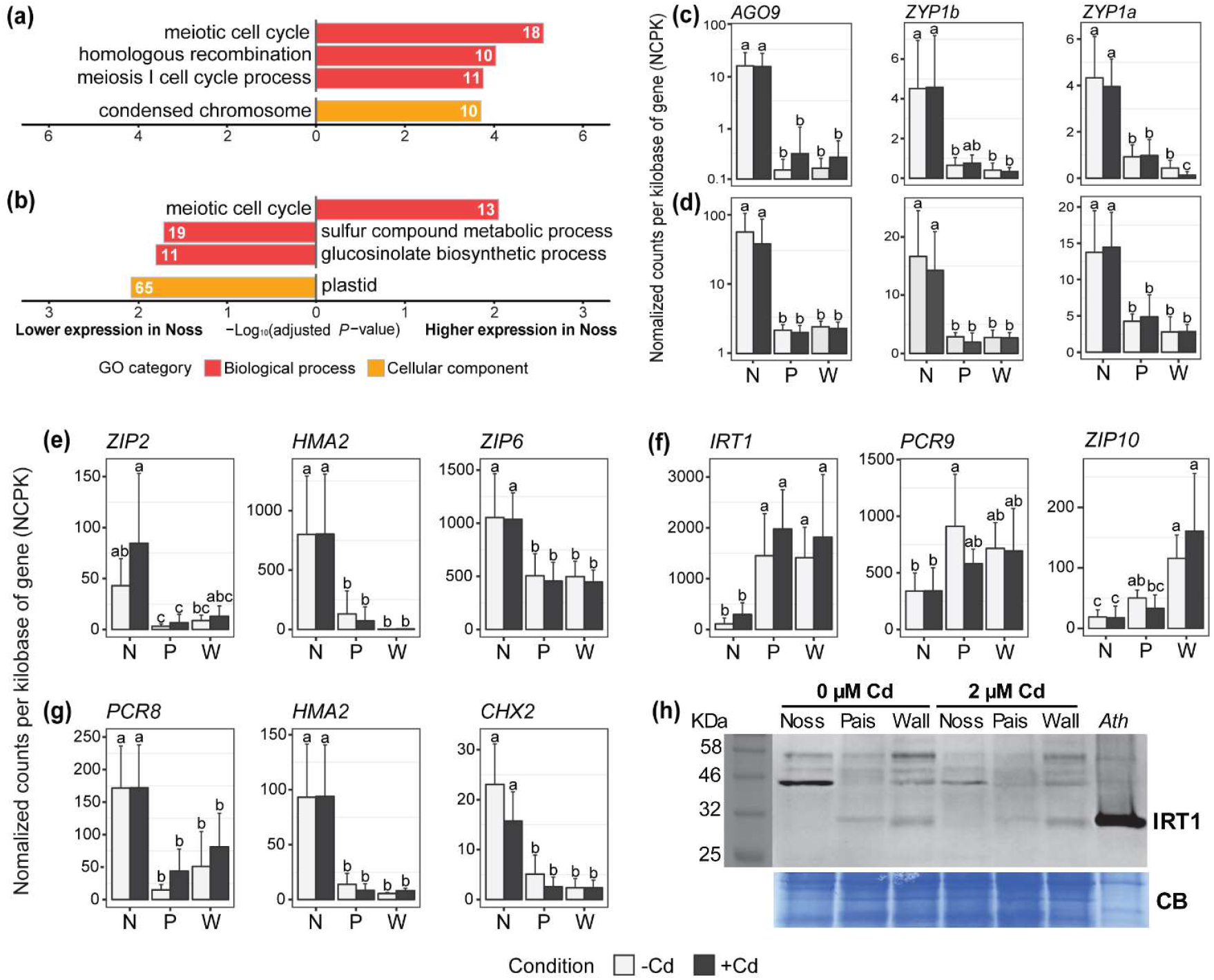
Candidate genes differentially expressed in Noss compared to both other populations Pais and Wall. (a, b) Significantly enriched gene ontology (GO) categories for roots (a) and shoots (b) of plants cultivated in 0 µM Cd (see Figure 2c and d, Dataset S3). Genes differentially expressed between Noss from M soil and both Pais and Wall from NM soils were subjected to a GO term enrichment analysis. The number of genes in each over-represented category is shown inside bars (see Dataset S4 for all genes). (c, d) Relative transcript levels of the top three genes (largest between-population differences) taken from (a) and (b) in the over-represented GO category meiotic cell cycle, for roots (c) and shoots (d). (e-g) Relative transcript levels of metal homeostasis genes of which transcript levels are high (e, g) and low (f) in Noss (N) compared to both Pais (P) and Wall (W), for roots (e, f) and shoots (g) (top three genes from Table S8). Bars show means ± SD (*n* = 6) of normalized counts, which were additionally normalized per kilobase of gene length (NCPK). Different characters represent statistically significant groups based on two-way ANOVA, followed by Tukey’s HSD test (*P <* 0.05, see Table S6, for details). Data are from the same experiment as shown in Figure 2. (H) Immunoblot of IRT1. Total protein extracts (40 µg) from roots of Noss_05, Pais_09 and Wall_07 were separated on a denaturing polyacrylamide gel, blotted, and detection was carried out using an anti-IRT1 antibody. Total protein extract (10 µg) from roots of Fe- and Zn-deficient *A. thaliana* (*Ath*, right lane) served as a positive control, with IRT1 detected as a single band at *ca*. 31 kDa. In *A. halleri*, additional bands at *ca*. 44, 48 and 56 kDa are likely to constitute non-specific ross-reactions of the antibody. Coomassie Blue (CB) stained membrane is shown as a loading control.

**Table 1.** List of genes in overrepresented meiosis- and homologous recombination-related GOs (see Fig. 3a, b).

| Name | Description | AGI | Aha Id. | GO<sup>a</sup> | Somatic DNA repair in *A. thaliana* | log<sub>2</sub>FC<sup>b</sup> |  |
| --- | --- | --- | --- | --- | --- | --- | --- |
|  |  |  |  |  |  | Max | Min |
| *AGO9* | Argonaute 9 | AT5G21150 | g12002 | Meiosis | Oliver *et al.*, 2014 | 6.57 *** | 3.88 *** |
| *ZYP1a* | Zipper-like 1a | AT1G22260 | g13785 | Meiosis/HR | n.i.<sup>c</sup> | 4.84 *** | 1.55 ** |
| *ZYP1b* | Zipper-like 1b | AT1G22275 | g13782 | Meiosis/HR | n.i. | 3.73 *** | 2.44 *** |
| *EMB2656* | Embryo defective 2656 | AT5G37630 | g20599 | HR | n.i. | 3.31 *** | 1.01 *** |
| *SMC2* | Structural maintenance of chromosomes 2 | AT3G47460 | g12815 | Meiosis | n.i. | 1.98 *** | 1.12 * |
| *SWI1* | SWITCH1 | AT5G51330 | g08416 | Meiosis | n.i. | 1.89 ** | 1.08 * |
| *ERCC1* | DNA excision repair protein ERCC-1 | AT3G05210 | g06284 | Meiosis | Dubest *et al.*, 2004 | 1.63 *** | 0.56 * |
| *MSH5* | MUTS-homologue 5 | AT3G20475 | g11121 | Meiosis/HR | n.i. | 1.60 *** | 1.04 ** |
| *PRD3* | Putative recombination initiation defects 3 | AT1G01690 | g01742 | Meiosis | n.i. | 1.60 *** | 1.05 * |
| *EME1A* | Essential meiotic endonuclease 1A | AT2G21800 | g23988 | Meiosis | Geuting *et al.*, 2009 | 1.59 *** | 0.72 * |
| *BRCA2B* | Breast cancer susceptibility 2 homolog B | AT5G01630 | g27928 | Meiosis/HR | Kumar *et al.*, 2019; Seeliger *et al.*, 2012 | 1.54 *** | -0.91 *** |
| NA | WEB family protein | AT3G02930 | g31838 | Meiosis/HR | n.i. | 1.47 *** | 0.55 *** |
| NA | Eisosome protein | AT1G04030 | g02033 | Meiosis | n.i. | 1.39 *** | 0.80 ** |
| *RPA2B* | Replication protein A, subunit RPA32 | AT3G02920 | g02469 | Meiosis/HR | Liu *et al.*, 2017 | 1.37 *** | 0.98 *** |
| *MHF2* | MPH1-associated histone-fold protein 2 | AT1G78790 | g01439 | Meiosis | n.i. | 1.32 *** | 0.58 * |
| *CAP-D3* | Condensin-2 complex subunit D3 | AT4G15890 | g20215 | Meiosis/HR | n.i. | 1.26 * | 0.94 * |
| *MCM8* | Minichromosome maintenance 8 | AT3G09660 | g06091 | Meiosis | n.i. | 1.21 ** | 0.82 * |
| *ATK1* | *Arabidopsis thaliana* knotted-like 1 | AT4G21270 | g29731 | Meiosis/HR | n.i. | 1.15 *** | 0.70 ** |
| *SMC3* | Structural maintenance of chromosome 3 | AT5G48600 | g15955 | Meiosis | n.i. | 1.13 *** | 0.46 * |
| *FLIP* | Fidgetin-like-1 interacting protein | AT1G04650 | g02098 | Meiosis | Fernandes *et al.*, 2018; Bouyer *et al.*, 2018 | 1.06 ** | 0.93 * |
| *HEB2* | Hypersensitive to excess boron 2 | AT3G16730 | g06531 | HR | Sakamoto *et al.*, 2011 | 0.88 *** | 0.40 * |
| *ESP* | Extra spindle poles | AT4G22970 | g14375 | Meiosis | n.i. | 0.82 ** | 0.42 * |
| *EME1B* | Essential meiotic endonuclease 1B | AT2G22140 | g28064 | Meiosis | Geuting *et al.*, 2009 | 0.70 *** | 0.35 ** |

<sup>a</sup>Meiosis, meiotic cell cycle or meiosis I cell cycle process; HR, homologous recombination (reciprocal meiotic recombination).
<sup>b</sup>Maximum and minimum of Log<sub>2</sub>(fold change Noss vs. any population from an NM site) across all tissues and conditions (adjusted *P*-value < 0.001, \*\*\*; < 0.01, \*\*; < 0.05, \*). <sup>c</sup>Not identified.

In the GO class meiotic cell cycle, transcript levels of the two gene copies of *ZIPper I* (*ZYP1a* and *ZYP1b*) were also substantially higher in Noss than in both Pais and Wall (Fig. 3e and f, Tables S5 and S6). Amino acid sequences of *At*ZYP1a and b are highly similar and conserved in *Ah*ZYP1a and b (Fig. S7). We validated sequencing-based transcriptomic data by reverse-transcription quantitative real-time PCR for a representative set of genes including *AGO9* and *ZYP1a/b* (Table S8, Fig. S9). An additional 15 genes in roots and 10 genes in shoots of the GO meiotic cell cycle showed elevated transcript levels in Noss compared to both populations of NM soil (see Fig. 3a and b, Dataset S4). The partly overlapping categories homologous recombination (GO:0035825), reciprocal meiotic recombination (GO:0007131), meiosis I cell cycle process (GO:0061982) and condensed chromosome (GO:0000793) were also enriched among transcripts of increased abundance in roots of Noss. Among a total of 23 genes that were more highly expressed in either roots or shoots of Noss compared to both Pais and Wall as well as grouped in over-represented GO terms related to meiotic cell cycle and homologous/meiotic recombination, homologs of eight genes (35%) were previously shown to have roles or implicated in somatic DNA repair in *A. thaliana* (Table 1). Furthermore, sulfur compound metabolic process (GO:0006790), glucosinolate biosynthetic process (GO:0019761) and plastid (GO:0009536) were enriched among transcripts of decreased abundance in shoot tissues of Noss (see Dataset S4).

### Differential abundance of metal homeostasis-related transcripts between populations

Next, we sought to identify metal homeostasis candidate genes contributing to the observed differences in Cd tolerance and accumulation between Noss and the two other populations. For this purpose, we intersected a manually curated list of metal homeostasis genes with the list of candidate transcripts (see Fig. 3e-g, |Log_2_FC| > 1, mean normalized counts across all samples > 2; Dataset S3, Table S7). Transcript levels of *Heavy Metal ATPase 2* (*HMA2*) and two *ZRT/IRT-like Protein* genes, *ZIP2* and *ZIP6*, were higher in roots of Noss than Pais and Wall (Fig. 3e). Root transcript levels of *Iron-Regulated Transporter 1* (*IRT1*), *Plant Cadmium Resistance 9* (*PCR9*) and *ZIP10* were lower in Noss (Fig. 3f). In shoots of Noss, transcript levels of *PCR8, HMA2* and *Cation/H*^*+*^ *Exchanger 2* (*CHX2*) were higher than in both Pais and Wall (Fig. 3g).

The IRT1 transmembrane transport protein is known as the high-affinity root iron (Fe^2+^) uptake system of dicotyledonous plants as well as the predominating path for inadvertent uptake of Cd^2+^ in Arabidopsis (Vert *et al*., 2002). Compared to Pais and Wall, an 84% to 92% lower expression of *IRT1* in Noss (see Fig. 3f) would provide an intuitive adaptive route towards plant Cd tolerance *via* exclusion (see Fig. 1d, Fig. S8). In *A. thaliana, IRT1* expression is under complex regulation at both the transcriptional and post-transcriptional levels (Kerkeb *et al*., 2008). To test whether reduced IRT1 protein levels accompany low *IRT1* transcript abundance in Noss, we carried out immunoblot detection using an anti-*At*IRT1 antibody in subsamples of root tissues from the same experiments. Note that amino acid sequences of IRT1 proteins of *A. halleri* Noss, Pais and Wall individuals as well as of *A. thaliana* are identical in the region corresponding to the peptide used to generate the anti-*At*IRT1 antibody (Fig. S8, boxed). At the size corresponding to IRT1 of Fe-deficient *A. thaliana* (Fig. 3h, right lane), we observed weak bands in both Pais and Wall cultivated under both control and 2 µM Cd exposure conditions, but no band was visible in Noss, in agreement with our observations at the transcript level (see also Fig. S9c for confirmation). Note that we loaded lower total protein from Fe-deficient *A. thaliana* in order not to overload the detection with a very strong signal and also because it served merely as a size marker. Note also that the extremely low levels of IRT1 protein under our non-Fe-deficient conditions can favor the detection of non-specific bands which do not appear when IRT1 protein levels are very high. Additional bands observed in *A. halleri* at higher molecular masses are likely to constitute non-specific cross-reactions of the antibody (44 kDa). The sizes of bands at approximately 48 and 56 kDa are also consistent with multiply ubiquitinated forms of IRT1, respectively, previously reported at distances of multiples of ∼9 kDa from IRT1 (Fig. 3h)(Barberon *et al*., 2011, Callis, 2014). In *A. thaliana*, the ubiquitination of IRT1 leads to its deactivation by removal from the plasma membrane through endocytosis (Barberon *et al*., 2011). Taking these data together, the absence of a signal corresponding to the size of functional IRT1 in Noss on immunoblots, as well as low IRT1 signals in Pais and Wall, with no changes upon cultivation in 2 µM Cd, were in full agreement with the between-population differences observed for *IRT1* transcript levels using RNA-seq and RT-qPCR (see Fig. 3f, Fig. S9a).

### Cd exposure elicits transcriptional Fe deficiency responses in both populations from NM sites

In the less Cd-tolerant populations, upon exposure to Cd, we found differential transcript abundance for 210 genes (upregulated 53, downregulated 157) in shoots and of 428 genes (upregulated 143, downregulated 285) in roots of Pais. A similar number of 197 (upregulated 65, downregulated 132) in roots and 243 (upregulated 98, downregulated 145) in shoots responded to Cd in Wall (see Fig. 2a and b, Dataset S5). Of these, the responses of 44 genes in roots and of 84 genes in shoots were shared between both populations from NM soil, a larger number than expected by chance (*P* < 10^−50^, hypergeometric test). Transcriptomic Cd responses of both Pais and Wall were enriched in Fe deficiency responses (Fig. 4, Dataset S6). The associated enriched GO categories included cellular response to iron ion starvation (GO:0010106) among the upregulated transcripts, as well as intracellular sequestering of iron ion (GO:0006880) and ferroxidase activity (GO:0004322) among the downregulated transcripts in both roots and shoots.

**Figure 4.**
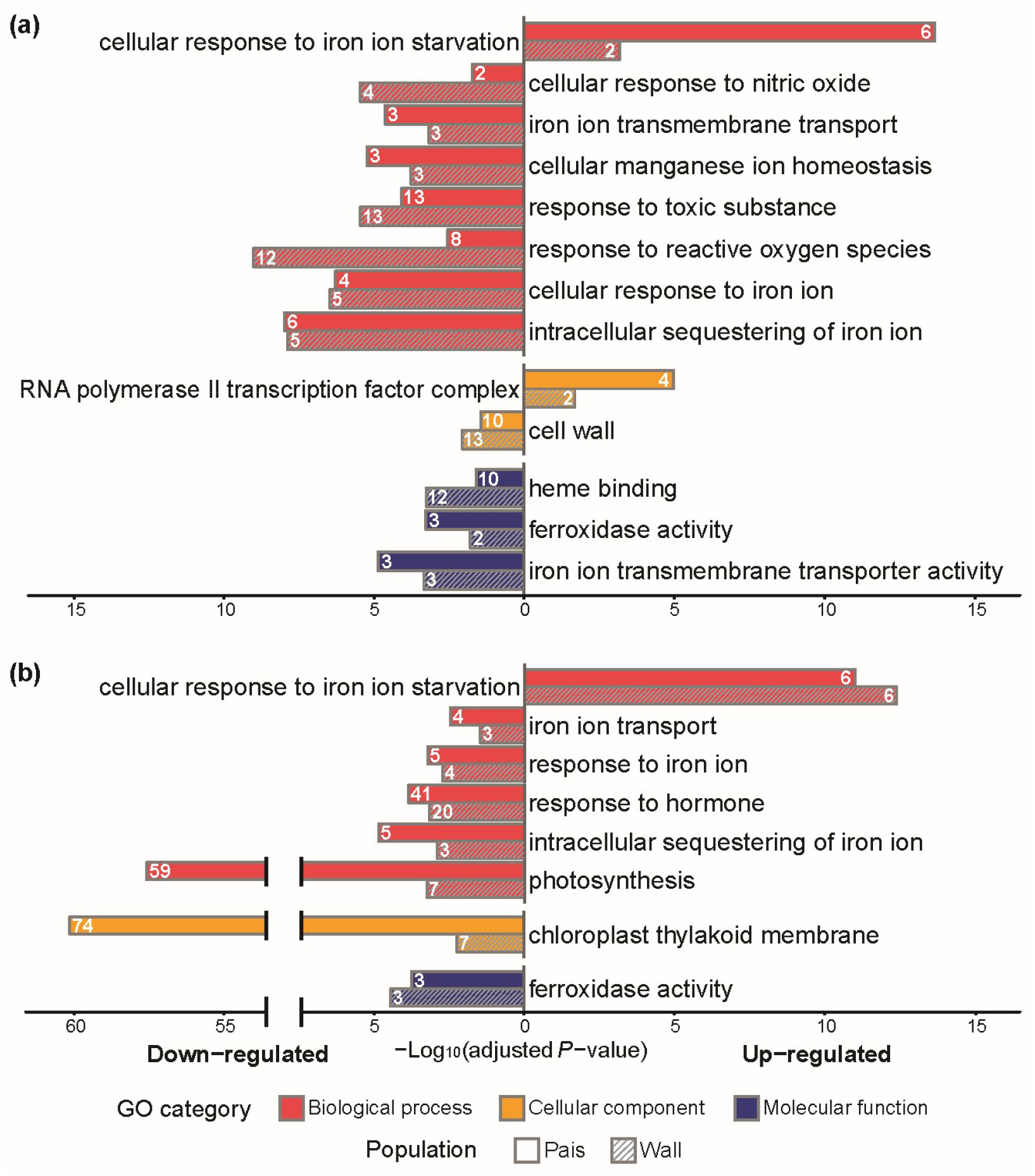
Functional classification of the transcriptional Cd responses in less metal-tolerant populations from NM sites. Significantly enriched GO categories among the transcriptional responses to Cd common in both the Pais and Wall populations for (a) roots and (b) shoots (see Figure 2a and b, Dataset S6). The numbers of genes in each over-represented category are shown inside bars. Data are from the same experiment as shown in Figure 2.

Among transcripts known to increase in abundance in *A. thaliana* under iron deficiency, transcript levels of *Popeye* (*PYE*), *Basic Helix-Loop-Helix 38/39/100/101* (*bHLH38/39/100/101*) transcription factors, *Brutus* (*BTS*) and *Brutus-like 1* (*BTSL1*), as well as *Ferric Reduction Oxidases* (*FRO1* and *FRO3*), *Nicotianamine Synthase 4* (*NAS4*), *Zinc Induced Facilitator 1* (*ZIF1*), *Oligopeptide Transporter 3* (*OPT3*) and *Natural Resistance-Associated Macrophage Protein 3* (*NRAMP3*) increased under Cd exposure (Table S9a). A number of genes have roles in iron storage/sequestration, and their transcript levels are downregulated under iron deficiency in *A. thaliana*. Of these, *Ferritin* (*FER1, FER3* and *FER4*) and *Vacuolar iron Transporter-Like protein* (*VTL1* and *VTL5*) transcript levels were downregulated in Cd-exposed Pais and Wall plants (Table S9b).

Further GOs enriched among transcripts downregulated in abundance in response to Cd were response to reactive oxygen species (GO:0000302), photosynthesis (GO:0015979), and chloroplast thylakoid membrane (GO:0009535). The gene content of these, however, is less compelling in implicating the respective biological processes (see Table S9b; Datasets S5 and S6). Chloroplast thylakoid membrane is a child term of plastid (GO:0009536), which was enriched among transcripts present at lower levels in Noss than in both Pais and Wall. Six genes were in common between these (Table S10), of which only one, *Light Harvesting Complex of photosystem II* (*LHCB4*.*2*), appears to function directly in photosynthesis. A more extensive search for transcripts that are Cd-responsive in both Pais and Wall as well as differentially expressed in the Noss population identified *Bifunctional inhibitor/lipid-transfer protein/seed storage 2S albumin superfamily protein* in the root and *NAD(P)-binding Rossmann-fold superfamily protein* in the shoot (Table S11). Finally, metal homeostasis protein-encoding transcripts upregulated in abundance upon Cd exposure in Pais or Wall were generally less abundant in Noss regardless of Cd treatment (Fig. S10, cluster R5 and S5). Conversely, metal homeostasis protein-coding transcripts that were decreased in abundance under Cd exposure in Pais or Wall were generally present at higher levels in Noss (Fig. S10, cluster R2 and S2). These between-population differences primarily comprised genes associated with Fe deficiency responses of *A. thaliana* as described above (see Fig. 4, Table S9), thus indicating against a constitutive proactive activation in more Cd-tolerant Noss of the transcriptional Cd responses observed here in the less Cd-tolerant Pais and Wall populations. These results suggested that transcriptional symptoms elicited by Cd exposure were observed already at far-subtoxic Cd concentrations, which neither caused growth impairment nor chlorosis, in the two less Cd-tolerant populations originating from NM soils.

### Lower transcript levels of long terminal repeat retrotransposons in Noss

Comparative transcriptomics suggested that in particular, *AGO9* transcript levels were higher in Noss compared to both Pais and Wall, with the largest quantitative difference within its functional context (see Table 1, Fig. 3c and d). If this were indeed the case, then we would expect that long terminal repeat retrotransposons (LTR-TEs), target loci known to be transcriptionally silenced dependent on *AGO9* in *A. thaliana*, are less transcriptionally active in *A. halleri* from Noss than *A. halleri* from Pais and Wall (Duran-Figueroa and Vielle-Calzada, 2010). In Noss, transcript levels derived from LTR-TEs were indeed lower in shoot tissues by comparison to both Pais and Wall and also lower in root tissues than in Pais (Fig. 5, Fig. S11, Dataset S7).

**Figure 5.**
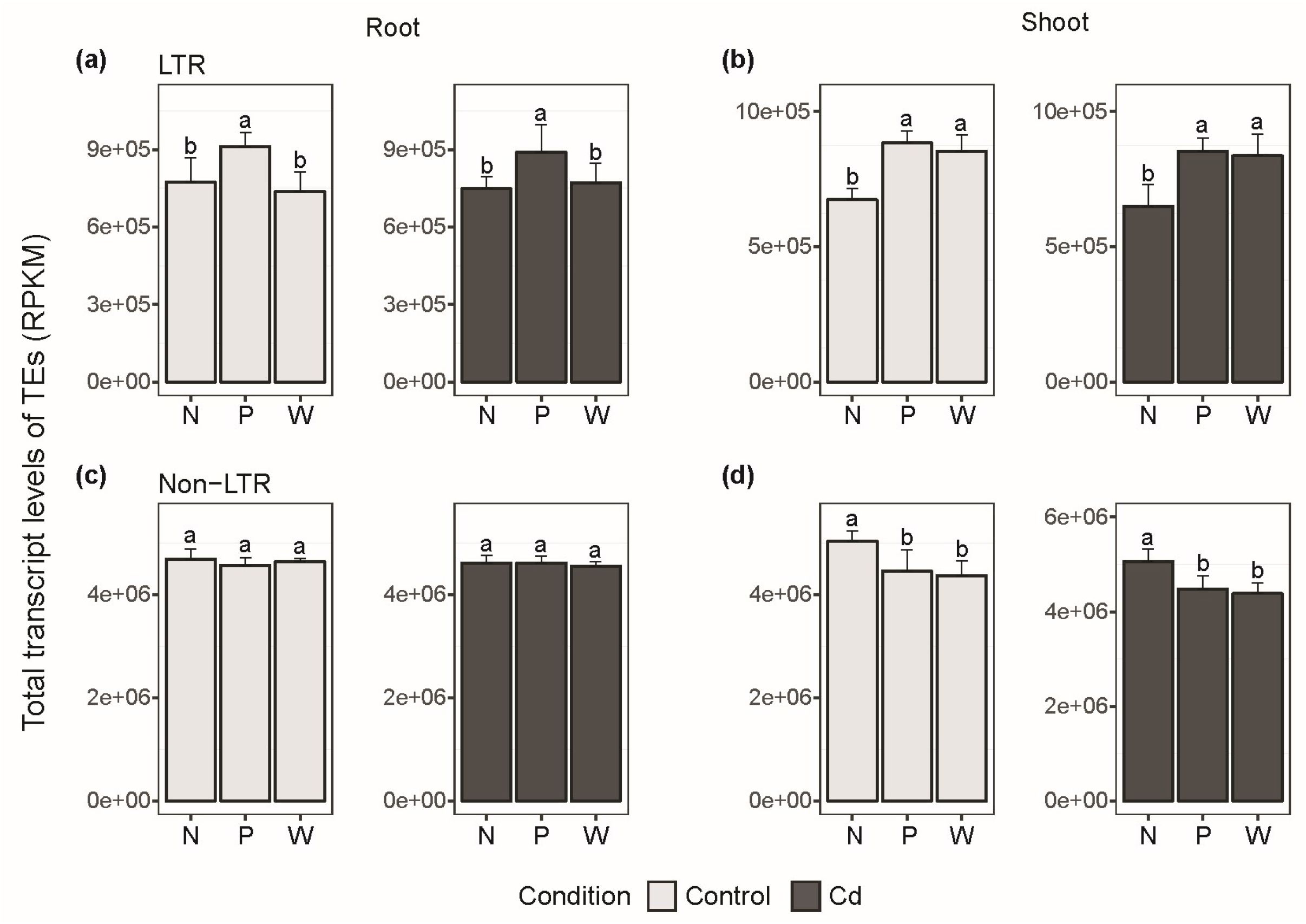
Genome-wide sum of transcript levels contributed by different types of transposable elements in Noss, Pais and Wall. (a-d) Bars show means ± SD (*n* = 6) of transcript levels of transposable elements (TEs), corresponding to the sum total of RPKM for LTR (Long terminal repeat) retrotransposons (a, b) and non-LTR TEs (c, d) in root (a, c) and shoot (b, d), with RPKM ≥ 7 per locus. Different characters represent statistically significant differences based on one-way ANOVA, followed by Tukey’s HSD test (*P <* 0.05).

### Elevated *AGO9* transcript levels in *A. halleri* originating from European M sites

In *A. halleri*, M sites were apparently colonized by individuals from neighboring NM sites (Pauwels *et al*., 2005). We identified two additional *A. halleri* population pairs combining a highly metal-contaminated (M) site and a non-contaminated (NM) site in a joint geographic region (Stein *et al*., 2017). For each of the three M-NM population pairs, clones of two individuals available in our laboratory (Stein *et al*., 2017) were cultivated hydroponically, and *AGO9* transcript levels were quantified in leaves by RT-qPCR. Within each of the three pairs, *AGO9* transcript levels were significantly higher in *A. halleri* of M soil origin than of NM soil origin (Fig. 6; see Fig. 3d, Dataset S2). Among individuals from M sites, there was strong quantitative variation in *AGO9* transcript abundance, but all means were higher than the means of individuals from the corresponding NM site of the same population pair. Taking into account the spatial heterogeneity of M habitats and the genetic diversity within outcrossing *A. halleri* (Krämer, 2018), we conclude that elevated expression of *AGO9* is common among individuals from M site populations in several regions of Europe.

**Figure 6.**
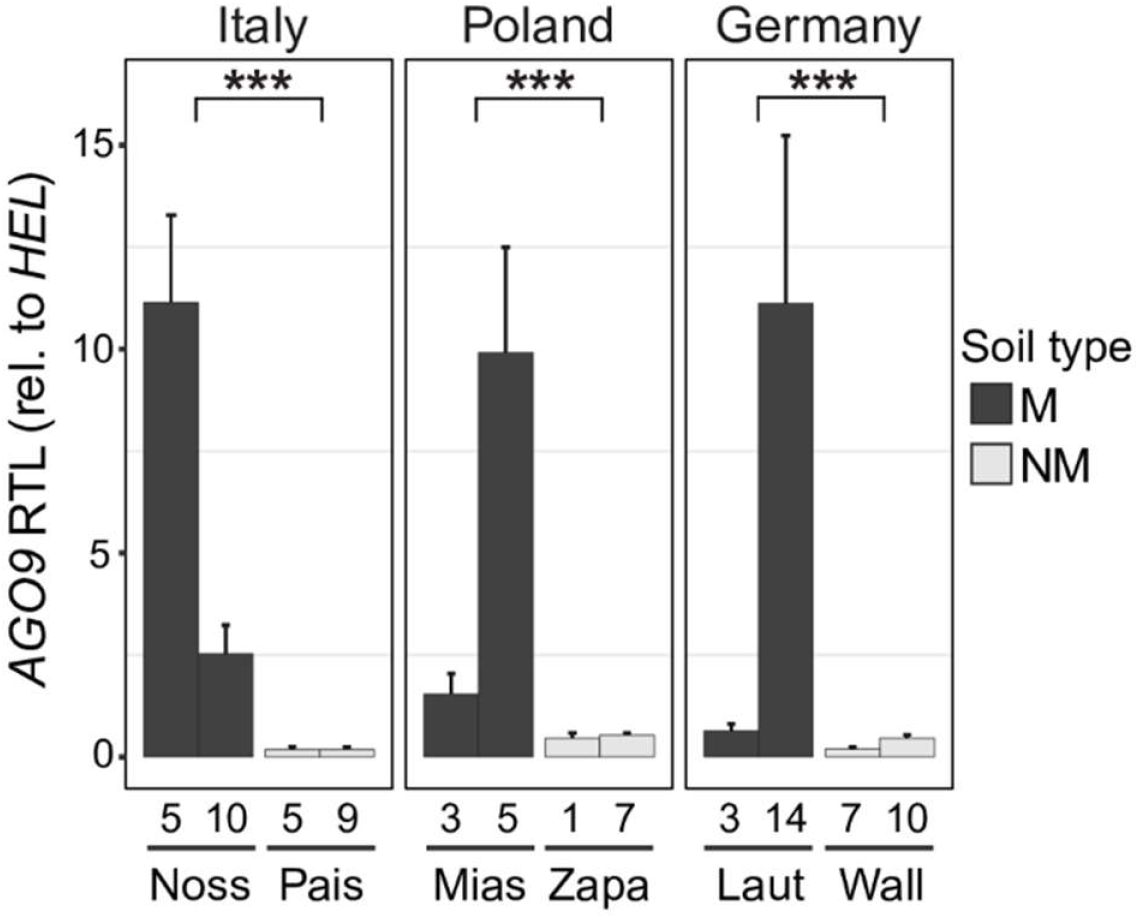
*ARGONAUTE 9* (AGO9) transcript levels in A. *halleri* individuals originating from geographically paired metalliferous and non-metalliferous sitess in Italy, Poland and Germany. Bargraphs show mean ± SD (*n* = 4) of relative *AGO9* transcript levels quantified by RT-qPCR in leaves of 5.5-week-old vegetative clones cultivated hydroponically (two independent PCR runs on each of two independently synthesized cDNAs from a homogenized pool of six replicate clones; all PCR reactions were conducted in hexuplicate). Data are shown relative to the constitutively expressed gene *Helicase* (*HEL*). Asterisks represent statistically significant differences between M and NM site of each pair, based on one-way ANOVA of Log-transformed data, followed by Tukey’s HSD test (***, *P <* 0.001). For details on collection sites and individuals see Stein et al. (2017).

## Discussion

### Divergent transcript levels between populations arise more abundantly than divergent transcriptomic responses to Cd

To globally identify molecular alterations associated with plant adaptation to extreme abiotic stress, we conducted comparative transcriptomics of *A. halleri* populations from contrasting environments of origin (Fig. 2). In line with their origin from the most highly Cd-contaminated *A. halleri* site known in Europe, our results support environment-dependent phenotypic differentiation in *A. halleri* and adaptation to extremely high soil Cd levels at Noss (Fig. 1). Transcript levels of a few hundred genes differed between Cd-exposed and unexposed *A. halleri* plants from the two populations originating from NM soils, Pais and Wall. Thus, these less Cd-tolerant plants attained a different transcriptomic state in the presence of Cd through a process that must involve the initial interaction of Cd^2+^ with primary Cd-sensing or -sensitive sites, of which the molecular identities are yet unknown. By contrast, in *A. halleri* from the more Cd-tolerant Noss (M) population, there was no single gene of which transcript levels differed between the Cd and the control treatment (Fig. 2). This implies that in Noss, Cd^2+^ was either effectively kept away from all Cd-sensing and -sensitive sites (Krämer, 2018), which is likely, or alternatively Cd^2+^ did not result in any transcriptomic adjustment within a timeframe relevant for our harvest after 16 d of exposure. Given that Noss is more Cd-tolerant than both Pais and Wall (see Fig. 1), the molecular mechanisms conferring tolerance must then either be active also in the absence of Cd, or alternatively their implementation must operate entirely downstream of transcript levels, for example at the translational or post-translational level.

Several thousand transcripts differed in abundance between the Noss (M) population of *A. halleri* and either one of the two populations originating from NM sites irrespective of Cd exposure (Fig. 2; see also Fig. S10). This suggests that between-population transcriptomic divergence is substantial even among the geographically proximal and phylogenetically closely related *A. halleri* populations Noss and Pais (Figs. S3, S11; Anderson *et al*., manuscript in preparation), by comparison to between-population differences in the responses to Cd exposure. Similarly, earlier cross-species comparative transcriptomics studies identified responses to metal exposure in *A. thaliana*, a species with only basal metal tolerance, whereas there were no remarkable additional species-specific responses among metal homeostasis genes in the metal-hypertolerant species *A. halleri* (Talke *et al*., 2006, Weber *et al*., 2006). Instead, tens of genes with predicted roles in metal homeostasis were more highly expressed in *A. halleri* compared to *A. thaliana* irrespective of the growth conditions.

### Constitutive global activation of meiosis-related genes

Meiosis is essential for sexual reproduction in eukaryotes and occurs exclusively in male and female gametophyte precursor cells during the reproductive phase of the life cycle (Mercier *et al*., 2015). Yet, in somatic tissues we observed an overrepresentation of the Gene Ontology Term (GO) “meiotic cell cycle” among transcripts of higher abundance in Noss by comparison to both Pais and Wall (Fig. 3, Table 1, Dataset S4). This included genes functioning during entry into meiosis (*AGO9, SWI1*), cohesion complex (*SMC3*), synaptonemal complex (*ZYP1a/b*), recombination and its control (*PRD3, BRCA2B, MHF2, MSH5, FLIP*), for example, in *A. thaliana* (Mercier *et al*., 2015). In addition to its role during meiosis, DNA double-strand break repair (DSBR) through homologous recombination (HR) functions to maintain genome integrity following DNA damage in somatic cells (Schuermann *et al*., 2005). Exposure of plants to an excess of aluminum, boron or heavy metal ions can cause DNA damage (Kovalchuk *et al*., 2001, Rounds and Larsen, 2008, Sakamoto *et al*., 2011, Morales *et al*., 2016). Of the meiosis-related genes identified here, there is evidence for an additional involvement in somatic HR-mediated DSBR for *BRCA2* (Seeliger *et al*., 2012), *EME1A/B* (Geuting *et al*., 2009), *HEB2* (Sakamoto *et al*., 2011), *FLIP* (Bouyer *et al*., 2018, Fernandes *et al*., 2018), *RPA2B* (Liu *et al*., 2017) and *ERCC1* (Dubest *et al*., 2004) in *A. thaliana* (Table 1; Dataset S4). *ERCC1* was also implicated in non-HR nucleotide excision repair (Hefner *et al*., 2003).

The two most strongly differentially expressed meiosis-related genes, *AGO9* and *ZYP1a/b*, were so far thought to function exclusively in reproductive development of *A. thaliana. AGO9* is required pre-meiotically to prevent the formation of excessive gametophytic precursor cells and expressed in a single layer of somatic companion cells surrounding the megaspore mother cell, as well as in pollen in *A. thaliana* (Olmedo-Monfil *et al*., 2010). *At*AGO9 was proposed to act in the silencing of pericentromeric TEs, predominantly long terminal repeat retrotransposons (LTR-TEs), in the ovule through the interaction with 24-nucleotide small RNAs (Duran-Figueroa and Vielle-Calzada, 2010). Like other organisms, plants generally inactivate TEs in order to maintain genomic stability. Exposure to abiotic stress, including also heavy metal exposure, can result in the expression and transposition of some TEs, especially LTR-TEs, in plants (Grandbastien, 2015) and other organisms (Horvath *et al*., 2017). In *A. halleri* from Noss elevated expression of *AGO9* in somatic cells may serve to counteract LTR-TE activation as an adaptation to their high-stress natural environment. Consistent with this hypothesis, transcripts derived from LTR-TEs were present at overall lower levels in both root and shoot tissues of Noss compared to Pais (Fig. 5).

We detected no global increase in the levels of transcripts derived from LTR-TEs or any other TEs at 2 µM Cd compared to control conditions. It is possible that an increase in TE expression is locus-specific, or occurs at different times after the onset of Cd exposure or at higher levels of Cd exposure which plants might encounter at the Noss site at least temporarily (Krämer, 2018), or in response to other abiotic stresses or combined stress exposure, in nature (see Table S1). Published circumstantial evidence has implicated *AGO9* in DNA damage repair in seedlings of *A. thaliana*, i.e. at the vegetative stage (Oliver *et al*., 2014). Available antibodies against *A. thaliana* AGO9 and ZYP1 did not detect specific bands in immunoblots of floral buds in our hands, as established by comparing wild-type Arabidopsis with the corresponding mutants (data not shown). Future work will address the biological functions of *AGO9* in somatic tissues of *A. halleri*.

We propose that our results reflect a constitutive transcriptional activation of genome integrity maintenance in vegetative tissues as a component of extreme abiotic stress adaptation of *A. halleri* at Noss. The genes newly implicated here in somatic genome integrity maintenance (Table 1) may have remained unidentified in the past because their functions in somatic cells are specific to *A. halleri* or because extremely low general expression levels may be sufficient to accomplish a possible somatic DNA repair function in *A. thaliana*. It should be noted that Cd is not the only abiotic stress factor at the Noss site in the field, where soil concentrations of Zn, Pb and Cu are also extremely high (Stein *et al*., 2017).

Of the meiosis genes showing elevated transcript levels in *A. halleri* from Noss (see Table 1), the four genes *ZYP1a/b, PRD3* and *SMC3* were among a total of eight meiosis genes identified to be under selection in autotretraploid *Arabidopsis arenosa*, for which there is no information on transcript levels to date (Yant *et al*., 2013). The predicted amino acid substitutions reported as hallmarks of tetraploidy in *A. arenosa*, when also divergent from diploid *A. thaliana* (Col), were all absent in *A. halleri* individuals from both Noss and Pais (data not shown). Moreover, all known *A. halleri* are diploid, including all populations of this study (Anderson *et al*., manuscript in preparation). We conclude that the alterations in meiosis gene expression in plants adapted to an extreme environment reported here are unrelated to published work on the adaptation of meiosis to tetraploidy that was initially conducted in *A. arenosa* (Bomblies *et al*., 2015).

### Large alterations in only few metal homeostasis functions in *A. halleri* from Noss

Attenuated Cd accumulation in *A. halleri* from Noss (see Fig. 1) observed here in hydroponics is consistent with earlier results from the cultivation of Noss and Wall on an artificially Cd-contaminated soil (Stein *et al*., 2017) and with observations in hydroponic culture on accessions collected nearby (Meyer *et al*., 2015, Corso *et al*., 2018). Among metal homeostasis genes, we observed substantially lower *IRT1* expression in roots of Noss compared to both Pais and Wall at both the transcript (Fig. 3f) and protein levels (Fig. 3h, Fig. S9c). This can account for both enhanced Cd hypertolerance and attenuated Cd accumulation in *A. halleri* from Noss, considering that IRT1 is the primary route for non-specific uptake of toxic Cd^2+^ into *A. thaliana* (Vert *et al*., 2002). Taking our data together, comparably high tissue levels of iron (Fig. S2e) in Noss plants appear to arise independently of IRT1 (Fig. 3h), which warrants further study.

ZIP2 is another member of the same protein family as IRT1 and could mediate the influx of divalent metals into the cytosol (Shanmugam *et al*., 2013). In *A. thaliana*, transcript levels of *ZIP2* were decreased under Mn and Fe deficiency (Milner *et al*., 2013). Upon Cd exposure, concentrations of Mn and Fe were lowered in roots of the both populations from NM sites compared to Noss (Fig. S2d, e). Based on its enhanced transcript levels in Noss, *ZIP2* could be a potential candidate gene for a role in root Fe and Mn uptake in Noss (Fig. 3e). In the Wall and Pais populations originating from NM soils, Fe levels were comparable under both treatment conditions, which was probably a result of compensation through Fe deficiency responses, by contrast to Mn concentrations which were decreased under Cd exposure in both populations originating from NM sites.

Distinctly higher transcript levels of *HMA2* in both roots and shoots of Noss (Fig. 3e and g) when compared to both the Pais and the Wall populations suggest that enhanced cellular export of Cd contributes to enhanced Cd hypertolerance in this population. Alongside HMA4, the plasma membrane efflux pump HMA2 mediates root-to-shoot translocation of Zn in *A. thaliana* and can also transport chemically similar Cd^2+^ ions (Hussain *et al*., 2004). The *HMA2* transcript is present in roots of Zn-deficient *A. thaliana*, in particular (Sinclair *et al*., 2018). A possible role of *HMA2* in the adaptation of *A. halleri* to local soil composition in Noss must involve functional alterations compared to *A. thaliana*.

### Cd exposure induces Fe deficiency responses in both populations from NM sites

In the less Cd-tolerant Wall and Pais populations originating from NM soils, we observed Fe deficiency response-like transcriptomic changes upon exposure to 2 µM Cd for 16 d (Fig. 4; Table S9). Plants that are non-hypertolerant to metals, for example *A. thaliana*, activate aspects of Fe deficiency responses when exposed to an excess of heavy metals, for example Cd (Leskova *et al*., 2017). Previously published results in *A. thaliana* were from heavy metal treatment conditions causing a mix of possible toxicity symptoms and acclimation responses. By contrast, in this present study the Cd treatment condition was far below toxicity thresholds of *A. halleri* that is known for species-wide Cd hypertolerance (see Fig. 1). Our data suggest that even under exposure to very low sub-toxic Cd concentrations, transcriptional responses of *A. halleri* reflect symptoms of nutrient imbalances caused by Cd. However, these responses to Cd are not associated with evolutionary adaptation to high-Cd soil in *A. halleri*.

A transcriptomics study reported Fe deficiency responses in *A. halleri* from the Polish M site PL22, but not in the I16 population from a metalliferous site close to Noss (M), in response to high-Zn exposure (Schvartzman *et al*., 2018). A highly Cd-hyperaccumulating accession of *Noccaea caerulescens* exhibited a Cd-induced Fe deficiency response, by contrast to a low Cd-accumulating accession (Halimaa *et al*., 2019). Both of these earlier studies were lacking the comparison with a neighboring population from an NM soil environment that is likely to reflect the most recent environment in evolutionary history, different from the work presented here. Consequently, central aspects of the heavy metal responses relating to Fe acquisition systems and tissue Fe contents according to this study cannot be directly compared to these earlier studies (Fig. S1, Table S9). A study that has just appeared online compares different *A. halleri* individuals from around Noss with *A. halleri* from a non-contaminated site around Lozio/IT (Stein *et al*., 2017, Corso *et al*., 2021). Corso et al. (2021) independently support between-population differential transcript levels of *IRT1, ZIP9, ZIP6, HMA2* and *FER4* (see Fig. 3 and S10). According to normalized transcript levels in the dataset of Corso et al. (2021), shoot *AGO9* and *ZYP1b* transcript levels were on average 1.7-fold and 4.6-fold higher, respectively, in *A. halleri* of M origin compared to NM origin.

In summary, this comparative study of *A. halleri* populations from different soil types of origin identifies transcriptomic alterations in plants that are adapted to an extreme environment, highly heavy metal-contaminated soil, which has arisen through rapid anthropogenic environmental change.

## Acknowledgements

We are grateful to Raphael Mercier (Max Planck Institute for Plant Breeding Research, Cologne, Germany) and Jean-Philippe Vielle-Calzada (CINVESTAV, Mexico City, Mexico) for sharing materials. We thank Petra Düchting, Andreas Aufermann and Jan Riering for assistance (Ruhr University Bochum, Germany). This work was supported by the Deutsche Forschungsgemeinschaft (Research Priority Program SPP1529 ADAPTOMICS, start-up grant to GL/JEA from grant number Kr1967/12 to UK, Research Priority Program SPP1819 RAPID EVOLUTION grant number Kr1967/16 to UK, and individual grant (Kr1967/3-3 to UK); and European Research Council Advanced Grant (grant number 788380 to UK). We acknowledge the Bielefeld-Giessen Center for Microbial Bioinformatics (BiGi) of the German Federal Ministry of Education and Research-funded German network for bioinformatics infrastructure (de.NBI, grant 031A533) for providing computational resources and related general support. Sequence data are available here: PRJEB35573, ENA, EMBL-EBI and MN747968-78 (submission ID: 2287280, Genbank, NCBI).

## Author Contribution

Designed research: UK, GL, JQ, JEA, BP; conducted experiments: GL, JQ, HA, VP, LS; analyzed data: GL, UK, JQ, HA, NJ, BP, LS; wrote paper: GL, UK, with contributions from JQ, HA, BP, VP.

## Figure Legends

**Fig. 1**. Cadmium tolerance and accumulation in three populations of *A. halleri*. (a) Proportion of individuals maintaining root growth (elongation) as a function of Cd concentration. Dashed vertical lines indicate ED_50_, the effective dose required to reach a proportion of 50% according to a dose-response model (see Table S2, Fig. S1). (b) Cd tolerance index. Bars represent mean ± SD of EC_100_ (effective Cd concentration causing 100% root growth inhibition) in a given plant individual (Schat and Ten Bookum, 1992). Plants were sequentially exposed to stepwise increasing concentrations of CdSO_4_ in hydroponic culture every week (*n* = 6 to 9 per population, with one to three vegetative clones of each three to six genotypes per population; (a, b)). (c) Cd accumulation efficiency at the site of origin calculated from published field survey data (bars represent means ± SD of *n* = 11 to 12 individuals in the field; units employed, µg Cd g^-1^ dry biomass in leaves, µg total Cd g^-1^ dry mass in soil; data from Stein et al. 2017). (d) Cd concentration in root and shoot tissues of hydroponically cultivated plants. Shown are mean ± SD (*n* = 12 to 20 clones per population comprising both genotypes, from all three replicate experiments; see Table S1). Four-week-old vegetative clones were exposed to 0 (-Cd) and 2 µM CdSO_4_ (+Cd) in hydroponic culture for 16 d alongside the plants cultivated for RNA-seq. Different characters denote statistically significant differences between means based on two-way ANOVA, followed by Tukey’s HSD test (Log-transformed data were used in (c)) (*P* < 0.05; see Table S3, for nested ANOVA of genotypes within a population (D)).

**Fig. 2**. Between-population differences according to transcriptome sequencing. (a, b) Venn diagrams show the numbers of genes exhibiting differential transcript abundances under exposure to Cd (+Cd) compared to control conditions (-Cd) for the three populations Noss (N), Pais (P) and Wall (W) in root (a) and shoot (b) tissues. Upregulation (red): Log_2_(fold change +Cd *vs*. –Cd) = Log_2_FC > 0.5; downregulation (blue): Log_2_FC < -0.5; all with adjusted *P*-value < 0.05. (c, d) Venn diagrams show the numbers of genes exhibiting differential transcript abundance between populations in root (c) and shoot (d) tissues of plants cultivated under Cd exposure (violet) or control conditions (orange) (|Log_2_FC| > 0.5; adjusted *P*-value < 0.05). Red lines surround genes differentially expressed between Noss and both populations of NM soils (Pais and Wall). Four-week-old vegetative clones were exposed to 2 µM Cd or 0 µM Cd (controls) in hydroponic culture for 16 d before harvest. Data are from three independent experiments (repeats), with two genotypes per population and material pooled from six replicate vegetative clones (three hydroponic culture vessels) per genotype in each experiment.

**Fig. 3**. Candidate genes differentially expressed in Noss compared to both other populations Pais and Wall. (a, b) Significantly enriched gene ontology (GO) categories for roots (a) and shoots (b) of plants cultivated in 0 µM Cd (see Fig. 2c and d, Dataset S3). Genes differentially expressed between Noss from M soil and both Pais and Wall from NM soils were subjected to a GO term enrichment analysis. The number of genes in each over-represented category is shown inside bars (see Dataset S4 for all genes). (c, d) Relative transcript levels of the top three genes (largest between-population differences) taken from (a) and (b) in the over-represented GO category meiotic cell cycle, for roots (c) and shoots (d). (e-g) Relative transcript levels of metal homeostasis genes of which transcript levels are high (e, g) and low (f) in Noss (N) compared to both Pais (P) and Wall (W), for roots (e, f) and shoots (g) (top three genes from Table S7). Bars show means ± SD (*n* = 6) of normalized counts, which were additionally normalized per kilobase of gene length (NCPK). Different characters represent statistically significant groups based on two-way ANOVA, followed by Tukey’s HSD test (*P <* 0.05, see Table S5, for details). Data are from the same experiment as shown in Fig. 2. (h) Immunoblot of IRT1. Total protein extracts (40 µg) from roots of Noss_05, Pais_09 and Wall_07 were separated on a denaturing polyacrylamide gel, blotted, and detection was carried out using an anti-IRT1 antibody. Total protein extract (10 µg) from roots of Fe- and Zn-deficient *A. thaliana* (*Ath*, right lane) served as a positive control, with IRT1 detected as a single band at *ca*. 31 kDa. In *A. halleri*, additional bands at *ca*. 44, 48 and 56 kDa are likely to constitute non-specific ross-reactions of the antibody. Coomassie Blue (CB) stained membrane is shown as a loading control.

**Fig. 4**. Functional classification of the transcriptional Cd responses in less metal-tolerant populations from NM sites. Significantly enriched GO categories among the transcriptional responses to Cd common in both the Pais and Wall populations for roots (a) and shoots (b) (see Fig. 2a and b, Dataset S6). The numbers of genes in each over-represented category are shown inside bars. Data are from the same experiment as shown in Fig. 2.

**Fig. 5**. Genome-wide sum of transcript levels contributed by different types of transposable elements in Noss, Pais and Wall. (a-d) Bars show means ± SD (*n* = 6) of transcript levels of transposable elements (TEs), corresponding to the sum total of RPKM for LTR (Long terminal repeat) retrotransposons (a, b) and non-LTR TEs (c, d) in root (a, c) and shoot (b, d), with RPKM ≥ 7 per locus. Different characters represent statistically significant differences based on one-way ANOVA, followed by Tukey’s HSD test (*P <* 0.05).

**Figure 6**. *ARGONAUTE 9* (*AGO9*) transcript levels in *A. halleri* individuals originating from geographically paired metalliferous and non-metalliferous sites in Italy, Poland and Germany. Bargraphs show mean ± SD (*n* = 4) of relative *AGO9* transcript levels quantified by RT-qPCR in leaves of 5.5-week-old vegetative clones cultivated hydroponically (two independent PCR runs on each of two independently synthesized cDNAs from a homogenized pool of six replicate clones; all PCR reactions were conducted in hexuplicate). Data are shown relative to the constitutively expressed gene *Helicase* (*HEL*). Asterisks represent statistically significant differences between M and NM site of each pair, based on one-way ANOVA of Log-transformed data, followed by Tukey’s HSD test (***, *P <* 0.001). For details on collection sites and individuals see Stein et al. (2017).

