## Supplemental Tables, Figures and Methods for "Constitutively enhanced genome integrity maintenance and direct stress mitigation characterize transcriptome of extreme stress-adapted *Arabidopsis halleri*"

**Original Research Article**

**This file includes:**

Supplemental Tables S1 to S11

Supplemental Figs. S1 to S11

Supplemental Experimental Procedures

Supplemental References

### Supplemental Tables

**Table S1.** Information on the field sites of origin of the three *A. halleri* populations in this study.

| Population name | Short name | Soil type | GPS coordinates | Population size | Individual Id. | Total concentration in soil (mg kg$^{-1}$) | | Extractable concentration in soil (mg kg$^{-1}$)[a] | | Concentration in leaves (mg kg$^{-1}$) | |
|---|---|---|---|---|---|---|---|---|---|---|---|
| | | | | | | Cd | Zn | Cd | Zn | Cd | Zn |
| Ponte Nossa (Italy) | Noss | M[1] | N45°51'33.5'' E09°52'36.1'' | 500-5000 | Noss_05 | 980 | 130,000 | 460 | 45,000 | 290 | 11,000 |
| | | | | | Noss_10 | 880 | 150,000 | 70 | 28,000 | 110 | 25,000 |
| Paisco Loveno (Italy) | Pais | NM[2] | N46°03'20.0'' E10°14'34.7'' | 500-5000 | Pais_05 | 1.4 | 110 | 0 | 20 | 5.6 | 4,000 |
| | | | | | Pais_09 | 1.5 | 180 | 0 | 60 | 4.5 | 33,000 |
| Wallenfels (Germany) | Wall | NM | N50°24'40.9'' E11°33'19.1'' | 50-500 | Wall_07 | 2.0 | 160 | 0 | 10 | 27 | 12,000 |
| | | | | | Wall_10 | 1.9 | 140 | 1 | 30 | 37 | 5,500 |

[1]Metalliferous. [2]Non-metalliferous. [a]Extracted in 2N HCl. Data from Stein et al. (2017).

**Table S2.** Dose-response models for the effect of Cd on root growth in the three *A. halleri* populations Noss, Pais and Wall (see Fig. 1a, Fig. S1).

| **Population** | **Soil type** | **$ED_{20}$ (± SE)[1]** | **$ED_{50}$ (± SE)[1A]** | **$ED_{80}$ (± SE)[1B]** | **Slope (± SE)[2]** | ***n*** |
|---|---|---|---|---|---|---|
| Noss | M[3] | 256.4 (± 34.9)[a] | 329.8 (± 16.5)[a] | 424.0 (± 22.7)[a] | 5.51 (± 1.50)[a] | 9 |
| Pais | NM[4] | 152.0 (± 12.8)[b] | 183.0 (± 6.4)[b] | 220.4 (± 16.0)[b] | 7.46 (± 2.10)[a] | 9 |
| Wall | NM | 160.2 (± 12.5)[b] | 195.9 (± 7.1)[b] | 239.5 (± 12.6)[b] | 6.89 (± 1.41)[a] | 6 |

[1/1A/1B]Fit-based estimates of the Cd concentrations in hydroponic solution that fully inhibit root growth in 20, 50 and 80% of individuals of a population, respectively (based here on one to three vegetative clones of each three to six genotypes per population, *n*: replicate plants). [2]Slope of the dose-response curves reflecting the range of within-population phenotypic variation (Ritz *et al.*, 2015). [3]Metalliferous soil, [4]Non-metalliferous soil. Different characters ([a],[b]) denote statistically significant differences between means in one column ($P < 0.05$). Note the difference between ED and EC (see Fig. 1b).

**Table S3.** Summary statistics of nested ANOVA for Fig. 1d.

| | DF[1] | SSD[2] | MSD[3] | F[4] | *P*-value |
|---|---|---|---|---|---|
| **Root** | | | | | |
| Error:Genotype | | | | | |
| Population | 2 | 248638 | 124319 | 3.80 | 0.15 |
| Residuals | 3 | | | | |
| Error:Within | | | | | |
| Treatment | 1 | 1218125 | 1218125 | 90.20 | 1.04E-14 |
| Population:Treatment | 2 | 177862 | 88931 | 6.59 | 0.00226 |
| Residuals | 79 | 1066833 | 13504 | | |
| **Shoot** | | | | | |
| Error:Genotype | | | | | |
| Population | 2 | 56811 | 28405 | 3.30 | 0.175 |
| Residuals | 3 | 25860 | 8620 | | |
| Error:Within | | | | | |
| Treatment | 1 | 746796 | 746796 | 103.42 | 5.19E-16 |
| Population:Treatment | 2 | 73269 | 36634 | 5.07 | 0.00846 |
| Residuals | 79 | 570476 | 7221 | | |

[1]Degrees of freedom. [2]Sum of square of standard deviation. [3]Mean square of standard deviation. [4]F ratio.

**Table S4.** Metadata for transcriptome sequencing experiments.

| Experiment[1] | Condition | Population | Genotype | Soil of origin | Sample Id. per tissue | |
|---|---|---|---|---|---|---|
| | | | | | Shoot | Root |
| 1 | Control[2] | Noss | Noss_05 | M | P1 | P37 |
| | | | Noss_10 | M | P2 | P38 |
| | | Pais | Pais_05 | NM | P3 | P39 |
| | | | Pais_09 | NM | P4 | P40 |
| | | Wall | Wall_07 | NM | P5 | P41 |
| | | | Wall_10 | NM | P6 | P42 |
| | Cd[3] | Noss | Noss_05 | M | P7 | P43 |
| | | | Noss_10 | M | P8 | P44 |
| | | Pais | Pais_05 | NM | P9 | P45 |
| | | | Pais_09 | NM | P10 | P46 |
| | | Wall | Wall_07 | NM | P11 | P47 |
| | | | Wall_10 | NM | P12 | P48 |
| 2 | Control | Noss | Noss_05 | M | P13 | P49 |
| | | | Noss_10 | M | P14 | P50 |
| | | Pais | Pais_05 | NM | P15 | P51 |
| | | | Pais_09 | NM | P16 | P52 |
| | | Wall | Wall_07 | NM | P17 | P53 |
| | | | Wall_10 | NM | P18 | P54 |
| | Cd | Noss | Noss_05 | M | P19 | P55 |
| | | | Noss_10 | M | P20 | P56 |
| | | Pais | Pais_05 | NM | P21 | P57 |
| | | | Pais_09 | NM | P22 | P58 |
| | | Wall | Wall_07 | NM | P23 | P59 |
| | | | Wall_10 | NM | P24 | P60 |
| 3 | Control | Noss | Noss_05 | M | P25 | P61 |
| | | | Noss_10 | M | P26 | P62 |
| | | Pais | Pais_05 | NM | P27 | P63 |
| | | | Pais_09 | NM | P28 | P64 |
| | | Wall | Wall_07 | NM | P29 | P65 |
| | | | Wall_10 | NM | P30 | P66 |
| | Cd | Noss | Noss_05 | M | P31 | P67 |
| | | | Noss_10 | M | P32 | P68 |
| | | Pais | Pais_05 | NM | P33 | P69 |
| | | | Pais_09 | NM | P34 | P70 |
| | | Wall | Wall_07 | NM | P35 | P71 |
| | | | Wall_10 | NM | P36 | P72 |

[1]Independent experiments (repeats). [2]0 µM $CdSO_4$ (-Cd) treatment. [3]2 µM $CdSO_4$ (+Cd) treatment.

**Table S5.** Transformation of data for statistical tests in Fig. 3c-h.

| Tissue | Gene | Original | | | | Log-transformed | | | | Box-cox-transformed | | | |
|---|---|---|---|---|---|---|---|---|---|---|---|---|---|
| | | Normality (Shapiro-Wilk test) | | Equality of variances (Levene's test) | | Normality (Shapiro-Wilk test) | | Equality of variances (Levene's test) | | Normality (Shapiro-Wilk test) | | Equality of variances (Levene's test) | |
| | | W | *P* | F | *P* | W | *P* | F | *P* | W | *P* | F | *P* |
| Root | *AGO9* | 0.77 | 0.0000 | 4.29 | 0.0046 | **0.93** | **0.0320** | **0.39** | **0.8512** | NA | NA | NA | NA |
| | *ZYP1b* | 0.87 | 0.0005 | 5.33 | 0.0018 | **0.93** | **0.0219** | **1.48** | **0.2252** | 0.92 | 0.0141 | 1.53 | 0.2109 |
| | *ZYP1a* | 0.91 | 0.0068 | 2.57 | 0.0478 | 0.97 | 0.5333 | 15.95 | 0.0000 | **0.97** | **0.3896** | **3.93** | **0.0073** |
| | *ZIP2* | 0.77 | 0.0000 | 7.07 | 0.0002 | **0.95** | **0.1019** | **0.34** | **0.8832** | NA | NA | NA | NA |
| | *HMA2* | 0.74 | 0.0000 | 1.64 | 0.1800 | **0.94** | **0.0391** | **5.47** | **0.0011** | 0.94 | 0.0452 | 6.99 | 0.0002 |
| | *ZIP6* | **0.95** | **0.1065** | **1.36** | **0.2672** | 0.91 | 0.0053 | 0.85 | 0.5245 | 0.91 | 0.0048 | 1.64 | 0.1787 |
| | *IRT1* | 0.95 | 0.1443 | 3.06 | 0.0238 | 0.95 | 0.1166 | 1.64 | 0.1788 | **0.98** | **0.7385** | **1.14** | **0.3612** |
| | *PCR9* | **0.94** | **0.0708** | **1.76** | **0.1522** | 0.94 | 0.0440 | 1.08 | 0.3893 | 0.99 | 0.9955 | 1.08 | 0.3908 |
| | *ZIP10* | 0.85 | 0.0002 | 3.50 | 0.0130 | **0.98** | **0.6000** | **2.04** | **0.1010** | 0.98 | 0.6033 | 1.16 | 0.3503 |
| Shoot | *AGO9* | 0.76 | 0.0000 | 3.84 | 0.0083 | 0.96 | 0.1547 | 3.28 | 0.0177 | **0.96** | **0.1732** | **2.26** | **0.0740** |
| | *ZYP1b* | 0.87 | 0.0006 | 5.10 | 0.0017 | **0.95** | **0.0854** | **0.26** | **0.9320** | 0.95 | 0.0744 | 0.28 | 0.9199 |
| | *ZYP1a* | 0.92 | 0.0141 | 1.70 | 0.1647 | 0.90 | 0.0047 | 1.57 | 0.1988 | **0.95** | **0.0796** | **1.08** | **0.3938** |
| | *PCR8* | **0.95** | **0.0826** | **1.73** | **0.1575** | 0.98 | 0.8809 | 0.41 | 0.8403 | 0.97 | 0.5532 | 0.22 | 0.9491 |
| | *HMA2* | 0.87 | 0.0005 | 11.20 | 0.0000 | **0.98** | **0.6069** | **1.32** | **0.2807** | 0.97 | 0.4084 | 1.63 | 0.1824 |
| | *CHX2* | 0.91 | 0.0065 | 1.70 | 0.1660 | 0.95 | 0.1165 | 1.10 | 0.3809 | **0.96** | **0.2193** | **0.91** | **0.4858** |

Transformation types employed are highlighted in bold and were chosen based on both normality and equality of variances for each gene.

**Table S6.** Read coverage of genomic DNA of Noss and Pais genotypes in *AGO9* compared to in the genome and in adjacent regions.

| Region for coverage | Genotype | | | |
|---|---|---|---|---|
| | Noss_05 | Noss_10 | Pais_05 | Pais_09 |
| Genome-wide average[1] | 16.5 | 13.1 | 18.6 | 20.0 |
| *AGO9*[2] | 17.5 | 14.9 | 20.1 | 21.6 |
| Upstream[3] | 17.7 | 20.7 | 22.2 | 22.3 |
| Downstream[4] | 15.8 | 13.7 | 19.0 | 18.0 |

gDNA sequence data were obtained from genomic DNA using Illumina TruSeq.
[1]Genome-wide average coverage of Illumina short reads of genomic DNA mapped to the reference genome, *A. halleri* ssp. *gemmifera* (Matsumura) O'Kane & Al-Shehbaz accession Tada mine (W302); [2]Read coverage in *AGO9* genomic region (translational start to stop codon; 4,671 bp); [3]Read coverage in 5'-region upstream of the *AGO9* translational start codon (-4,671 bp to 0 bp); [4]Read coverage in 3'-region downstream of the *AGO9* translational stop codon (4,672 bp to 9,343 bp).

**Table S7.** Metal homeostasis genes for which transcript levels in Noss individuals (M soil origin) differ from both Pais and Wall individuals (NM soil of origin).

| Name | AGI | Aha Id. | Description | NCPK[b] (-Cd in Noss) Mean ± SD | $Log_2FC$ in -Cd[a] | |
|---|---|---|---|---|---|---|
| | | | | | Noss *vs*. Pais | Noss *vs*. Wall |
| **Root** | | | | | | |
| *ZIP2* | AT5G59520 | g11007 | ZRT/IRT-like protein 2 | 43 ± 27 | 3.8 *** | 2.3 ** |
| *HMA2* | AT4G30110 | g04213 | Heavy metal ATPase 2 | 800 ± 488 | 2.6 * | 8.0 *** |
| *ZIP6* | AT2G30080 | g24253 | ZRT/IRT-like protein 6 | 1,068 ± 440 | 1.1 ** | 1.1 ** |
| NA | AT5G21105 | g12006 | L-ascorbate oxidase, putative | 249 ± 77 | 1.0 *** | 1.1 *** |
| *ZIP10* | AT1G31260 | g27549 | ZRT/IRT-like protein 10 | 19 ± 12 | -1.4 * | -2.6 *** |
| *PCR9* | AT1G58320 | g24696 | Plant cadmium resistance 9 | 343 ± 156 | -1.4 ** | -1.1 * |
| *IRT1* | AT4G19690 | g15474 | Iron-regulated transporter 1 | 112.7 ± 112.7 | -3.7 *** | -3.7 *** |
| **Shoot** | | | | | | |
| *PCR8* | AT1G52200 | g15174 | Plant cadmium resistance 8 | 171 ± 63 | 3.5 *** | 1.8 ** |
| *HMA2* | AT4G30110 | g04213 | Heavy metal ATPase 2 | 92 ± 47 | 2.8 *** | 4.1 *** |
| *CHX2* | AT1G79400 | g01507 | Cation/$H^+$ exchanger 2 | 23 ± 8 | 2.2 *** | 3.3 *** |
| *YSL2* | AT5G24380 | g25310 | Yellow stripe like 2 | 43 ± 29 | 1.4 * | 1.4 * |

[a] $|Log_2(\text{fold change})| > 1$, adjusted $P < 0.05$, mean normalized counts across all samples > 2. To focus on the subset of the most relevant candidate transcripts, we employed a more stringent $Log_2FC$. Here this is additionally based on our expectation that the direct role in metal handling of the proteins encoded by this group of transcripts results in a strongly elevated quantitative requirement and thus comparably large differences. [b]NCPK: normalized counts per kilobase of gene length *$P < 0.05$, ** $P < 0.01$, *** $P < 0.001$.

**Table S8.** Primer pairs used for RT-qPCR.

| Gene | Strand | Length | Sequence | Primer efficiency (Mean) |
|---|---|---|---|---|
| *AGO9*[a] | F | 24 | GGCTTATCCCTGAATATTGACACT | 1.95 |
| | R | 22 | TGCAAGCAGGAAATCAACTACA | |
| *ZYP1a* | F | 23 | ACAAATCATGAAGGACAAGGAGC | 1.97 |
| | R | 23 | TCAGGTCATATTGCTTCTTTGCT | |
| *HMA2* | F | 24 | CCTCTCTCAGTTCCAAATCGTTAA | 1.92 |
| | R | 24 | GTTTCTCCGGTTACCCTCACATTT | |
| *ZIP6* | F | 23 | CTAGGGATGGTGATATTTGCAGC | 1.96 |
| | R | 24 | AGAATGATCCCAATAAACCCTCCA | |
| *IRT1* | F | 22 | CCTCCAGGCTGAGTATACGAAT | 1.98 |
| | R | 22 | TCCCTAACGCTATTCCGAATGG | |
| *EF1a* | F | 23 | TGAGCACGCTCTTCTTGCTTTCA | 2.03 |
| | R | 24 | GGTGGTGGCATCCATCTTGTTACA | |
| *AGO9*[b] | F | 21 | ACGAAGGAATGTTCAGGGAGC | |
| | R | 23 | ATGTGCTCTGGTTTCCTCTTCTC | |
| *HEL* | F | 21 | CCATTCTACTTTTTGGCGGCT | |
| | R | 23 | TCAATGGTGACTGATCCACTCTG | |

[a]Supplemental Material; [b]Fig. 6.

**Table S9.** Genes of which transcript levels responded to Cd in both Pais and Wall.

(a) Genes of which the levels were upregulated under Cd exposure relative to control conditions.

| Name | AGI | Aha Id.[3] | Description | NCPK (+Cd in Pais) Mean ± SD[4] | $Log_2$FC (+Cd *vs.* -Cd)[5] | |
|---|---|---|---|---|---|---|
| | | | | | Pais | Wall |
| **Root** | | | | | | |
| *FRO1*[1] | AT1G01590 | g01760 | Ferric reduction oxidase 1 | 110 ± 110 | 5.3 ***[6] | 3.7 *** |
| NA | NA | g19004 | Unknown protein | 760 ± 410 | 3.7 *** | 1.8 * |
| NA | NA | g19005 | Unknown protein | 290 ± 170 | 3.6 *** | 2.6 *** |
| NA | NA | g25721 | Unknown protein | 2,600 ± 1,100 | 3.4 *** | 2.8 *** |
| NA | NA | g02411 | Unknown protein | 190 ± 110 | 2.9 *** | 1.8 ** |
| *bHLH101*[1] | AT5G04150 | g15774 | Basic helix-loop-helix protein 101 | 390 ± 200 | 2.8 *** | 2.4 ** |
| NA | NA | g19003 | Unknown protein | 720± 290 | 2.7 *** | 2.2 *** |
| NA | NA | g19002 | Unknown protein | 1,300 ± 400 | 2.1 *** | 1.5 * |
| *bHLH39*[1] | AT3G56980 | g21154 | Basic helix-loop-helix protein 39 | 1,300 ± 500 | 2.1 *** | 1.9 ** |
| *NPF5.9* | AT3G01350 | g02612 | NRT1/ PTR family 5.9 | 410 ± 250 | 1.8 ** | 1.5 *** |
| *ORG1*[1] | AT5G53450 | g23050 | OBP3-responsive protein 1 | 810 ± 310 | 1.7 *** | 1.1 *** |
| **Shoot** | | | | | | |
| NA | NA | g17415 | Zinc transporter 7-like | 21 ± 22 | 6.3 *** | 6.0 *** |
| *bHLH39*[1] | AT3G56980 | g21154 | Basic helix-loop-helix protein 39 | 670 ± 250 | 3.9 *** | 3.6 *** |
| *bHLH38*[1] | AT3G56970 | g21155 | Basic helix-loop-helix protein 38 | 280 ± 120 | 3.8 *** | 3.5 *** |
| NA | NA | g19004 | Unknown protein | 780 ± 300 | 3.5 *** | 2.6 ** |
| NA | NA | g19005 | Unknown protein | 400 ± 220 | 3.4 *** | 2.9 *** |
| NA | AT1G12030 | g04876 | Phosphoenolpyruvate carboxylase, putative (DUF506) | 550 ± 280 | 3.3 *** | 2.2 *** |
| *bHLH100*[1] | AT2G41240 | g24495 | Basic helix-loop-helix protein 100 | 450 ± 270 | 3.3 * | 5.5 *** |
| NA | AT1G62420 | g17830 | DUF506 family protein | 380 ± 110 | 3.2 *** | 3.0 ** |
| NA | NA | g19002 | Unknown protein | 1,900 ± 700 | 3.0 *** | 2.6 *** |
| *MIOX4* | AT4G26260 | g10190 | Myo-inositol oxygenase 4 | 34 ± 58 | 2.8 * | 2.5 * |
| NA | AT2G14247 | g19693 | Expressed protein | 5,100 ± 1,500 | 2.6 ** | 2.4 *** |
| NA | NA | g19003 | Unknown protein | 3,500 ± 800 | 2.6 *** | 2.0 *** |
| NA*S4*[1] | AT1G56430 | g24765 | Nicotianamine synthase 4 | 410 ± 190 | 2.4 *** | 1.3 ** |
| *BTSL1*[1] | AT1G74770 | g01014 | BRUTUS like 1 | 44 ± 18 | 2.4 * | 2.7 * |
| *bZIP39* | AT2G36270 | g03527 | Basic-leucine zipper (bZIP) transcription factor 39 | 22 ± 12 | 2.2 *** | 1.8 * |
| *TSO2* | AT3G27060 | g20504 | Ribonucleoside-diphosphate reductase small chain C | 2,300 ± 700 | 2.2 *** | 1.1 *** |
| *WRKY12* | AT2G44745 | g28045 | WRKY family transcription factor 12 | 32 ± 13 | 2.2 ** | 1.3 ** |
| NA | AT5G05250 | g26041 | Hypothetical protein | 840 ± 230 | 2.1 *** | 1.6 ** |
| *NPF5.9* | AT3G01350 | g02612 | NRT1/ PTR family 5.9 | 390 ± 190 | 2.0 ** | 1.8 *** |
| NA | NA | g25721 | Unknown protein | 19,000 ± 8,000 | 2.0 *** | 1.3 * |
| *FRO1*[1] | AT1G01590 | g01760 | Ferric reduction oxidase 1 | 590 ± 320 | 1.9 *** | 2.0 *** |
| *FRO3*[1] | AT1G23020 | g13702 | Ferric reduction oxidase 3 | 720 ± 220 | 1.9 *** | 1.6 *** |
| NA | AT1G01570 | g01764 | Glycosyl group transferase | 5.5 ± 3.0 | 1.8 * | 1.7 *** |
| *ZIF1*[1] | AT5G13740 | g19325 | Zinc induced facilitator 1 | 1,100 ± 200 | 1.8 *** | 1.7 *** |
| *BTS*[1] | AT3G18290 | g21576 | Zinc finger protein BRUTUS | 2,700 ± 500 | 1.8 *** | 1.5 *** |
| *ORG1*[1] | AT5G53450 | g23050 | OBP3-responsive protein 1 | 2,400 ± 600 | 1.7 *** | 1.6 *** |
| *OPT3*[1] | AT4G16370 | g23102 | Oligopeptide transporter 3 | 3,900 ± 700 | 1.5 *** | 1.2 *** |
| *B1* | AT4G25700 | g10252 | Beta-hydroxylase 1 | 440 ± 130 | 1.3 *** | 1.7 *** |
| *PYE*[1] | AT3G47640 | g12843 | POPEYE | 430 ± 80 | 1.3 *** | 1.1 * |
| *NRAMP4*[1] | AT5G67330 | g07320 | Natural resistance-associated macrophage protein 4 | 3,100 ± 1,000 | 1.2 *** | 1.0 *** |
| *CGLD27*[1] | AT5G67370 | g07324 | Conserved in the green lineage and diatoms 27 | 3,600 ± 1,000 | 1.2 *** | 1.1 ** |

(b) Genes of which transcript levels were downregulated under Cd exposure relative to control conditions.

| Name | AGI | Aha Id.[3] | Description | NCPK (-Cd in Pais) Mean ± SD[4A] | $Log_2$FC (+Cd *vs.* -Cd)[5] | |
|---|---|---|---|---|---|---|
| | | | | | Pais | Wall |
| **Root** | | | | | | |
| *VTL5*[1A] | AT3G25190 | g30123 | Vacuolar iron transporter-like 5 | 67 ± 31 | -4.9 *** | -2.9 ** |
| *CYP82C4*[1] | AT4G31940 | g00781 | Cytochrome P450, family 82, subfamily C, polypeptide 4 | 360 ± 320 | -4.7 *** | -6.2 *** |
| NA | AT1G73120 | g04063 | F-box/RNI (RNase inhibitor) superfamily protein | 210 ± 120 | -4.3 *** | -4.3 *** |
| *VTL1*[1A] | AT1G21140 | g13452 | Vacuolar iron transporter-like 1 | 230 ± 80 | -3.1 *** | -3.2 *** |
| NA | NA | g00255 | Peroxidase superfamily protein | 1,400 ± 760 | -3.1 *** | -2.3 *** |
| *ELS1* | AT5G19700 | g12140 | Early leaf senescence 1 | 130 ± 54 | -2.8 *** | -1.9 ** |
| *IAN1* | AT1G33830 | g17730 | Immune-associated nucleotide-binding protein 1 | 11 ± 5.9 | -2.7 *** | -1.3 *** |
| NA | AT5G38940 | g24630 | RmlC-like cupins superfamily protein | 3,700 ± 1,300 | -2.6 *** | -2.2 ** |
| *ZIP9* | AT4G33020 | g32214 | ZRT/IRT-like protein 9 | 1,200 ± 260 | -2.3 *** | -1.6 * |
| NA | NA | g28793 | Protein kinase-like protein | 54 ± 34 | -2.2 ** | -1.6 * |
| *PER2* | AT1G05250 | g02157 | Peroxidase 2 | 4,900 ± 1,400 | -2.1 ** | -1.6 * |
| *CYP78A6* | AT2G46660 | g09658 | Cytochrome P450, family 78, subfamily A, polypeptide 6 | 98 ± 32 | -2.0 ** | -1.3 ** |
| *FER1*[1A, 2] | AT5G01600 | g27931 | Ferritin 1 | 1,900 ± 370 | -1.9 *** | -1.8 *** |
| *PER44* | AT4G26010 | g10217 | Peroxidase 44 | 1,400 ± 450 | -1.8 ** | -1.3 * |
| NA | NA | g02159 | Peroxidase superfamily protein | 1,900 ± 410 | -1.8 *** | -1.5 * |
| *FER3*[1A] | AT3G56090 | g11645 | Ferritin 3 | 88 ± 25 | -1.6 *** | -1.1 * |
| *DGR1* | AT1G80240 | g01629 | Duf642 L-GalL responsive gene 1 | 620 ± 350 | -1.5 ** | -1.8 ** |
| **Shoot** | | | | | | |
| *AED3* | AT1G09750 | g28855 | Apoplastic EDS1-dependent protein 3 | 510 ± 380 | -3.1 *** | -1.5 * |
| NA[1A] | AT1G68650 | g14724 | GDT1-like protein 5 | 73 ± 37 | -3.0 *** | -2.2 ** |
| NA | AT1G04540 | g02084 | Calcium-dependent lipid-binding family protein | 12 ± 61 | -2.8 *** | -2.0 * |
| *ENH1* | AT5G17170 | g00054 | Enhancer of sos3-1 | 1,200 ± 710 | -2.8 *** | -1.4 * |
| *FD2*[2] | AT1G60950 | g19915 | Ferredoxin 2 | 27,000 ± 14,000 | -2.4 *** | -1.1 * |
| *PSAN*[2] | AT5G64040 | g12421 | Photosystem I reaction center subunit PSI-N | 16,000 ± 8,800 | -2.4 *** | -1.1 * |
| *FER1*[1A] | AT5G01600 | g27931 | Ferritin 1 | 3,500 ± 2,400 | -2.4 *** | -1.9 *** |
| *FER4*[1A] | AT2G40300 | g12617 | Ferritin 4 | 330 ± 190 | -2.2 *** | -1.1 * |
| NA | NA | g05038 | Unknown protein | 74 ± 69 | -1.9 ** | -1.3 ** |
| NA | AT3G61870 | g05411 | Plant/protein | 2,300 ± 1,100 | -1.7 *** | -1.1 ** |
| *FTRA2*[2] | AT5G08410 | g30310 | Ferredoxin/thioredoxin reductase subunit A2 | 1,200 ± 510 | -1.7 *** | -1.1 * |
| *CHAT* | AT3G03480 | g26820 | Acetyl CoA:(Z)-3-hexen-1-ol acetyltransferase | 1,200 ± 1,700 | -1.7 *** | -1.3 * |
| *SFH3* | AT2G21540 | g23962 | SEC14-like 3 | 56 ± 31 | -1.6 ** | -1.3 * |
| *APE2* | AT4G36050 | g00297 | Apurinic-apyrimidinic endonuclease | 73 ± 180 | -1.6 *** | -1.4 * |
| *FER3*[1A, 2] | AT3G56090 | g11645 | Ferritin 3 | 380 ± 220 | -1.5 *** | -1.1 * |
| *APX1* | AT1G07890 | g02454 | Ascorbate peroxidase 1 | 2,800 ± 1,400 | -1.5 *** | -1.6 *** |
| *EXPB3* | AT4G28250 | g04426 | Expansin B3 | 740 ± 280 | -1.5 * | -1.5 ** |
| *YSL1*[1A] | AT4G24120 | g10414 | Yellow stripe like 1 | 240 ± 170 | -1.5 * | -1.2 * |
| *CXE8* | AT2G45600 | g09782 | Carboxylesterase 8 | 53 ± 230 | -1.5 *** | -1.5 *** |
| NA | AT3G61820 | g05417 | Aspartyl protease family protein | 340 ± 190 | -1.5 *** | -1.2 *** |
| *RTP1* | AT1G70260 | g03763 | Resistance to phytophthora parasitica 1 | 220 ± 120 | -1.4 ** | -1.4 ** |
| *FAX6* | AT3G20510 | g11131 | Fatty acid export 6 | 230 ± 77 | -1.4 *** | -1.5 * |
| *NHR2A* | AT5G45410 | g07576 | Non host resistance 2A | 1,100 ± 460 | -1.3 *** | -1.2 *** |
| *GLB3*[1A] | AT4G32690 | g00695 | Hemoglobin 3 | 75 ± 20 | -1.3 *** | -1.3 ** |
| *NHR2B* | AT4G25030 | g10315 | Non host resistance 2B | 480 ± 230 | -1.1 *** | -1.1 ** |

[1/1A]Transcript levels up-/down-regulated under Fe deficiency in *A. thaliana*, respectively. [2]Genes in overrepresented GO term "photosynthesis". [3]Gene ID from *Arabidopsis halleri* ssp. *gemmifera* genome assembly (Briskine *et al.*, 2017). [4/4A]Mean NCPK (normalized counts per kilobase of gene length) under +Cd/-Cd, respectively, in root (a) or shoot (b) of Pais. [5]|$Log_2$(fold change +Cd *vs.* -Cd)| > 1; see Table S7. [6]Adjusted *P*-value < 0.001, ***; < 0.01, **; < 0.05, *).

**Table S10.** Genes for all GO terms enriched among differentially expressed genes in Noss compared to the other two populations, which are also Cd-responsive in either Pais or Wall.

| | | | | Enriched GO[1] | | Log2FC | | | |
|---|---|---|---|---|---|---|---|---|---|
| **Name** | **AGI** | **Aha Id.** | **Description** | **GO term Id.** | **GO name** | **Noss *vs.* Pais (-Cd)** | **Noss *vs.* Wall (-Cd)** | **+Cd *vs.* -Cd in Pais** | **+Cd *vs.* -Cd in Wall** |
| **Root** | | | | | | | | | |
| No gene | | | | | | | | | |
| **Shoot** | | | | | | | | | |
| *NUDT24* | AT5G19470 | g13120 | Nudix hydrolase homolog 24 | GO:0009536 | plastid | -2.9 *** | -3.5 *** | -1.4 | -2.1 * |
| *TGG1* | AT5G26000 | g14035 | Thioglucoside glucohydrolase 1 | GO:0009536/GO:0006082 | plastid/sulfur compound metabolic process | -1.9 *** | -2.2 *** | -1.6 ** | -0.4 |
| NA | AT1G26090 | g08132 | P-loop containing nucleoside triphosphate hydrolases superfamily protein | GO:0009536 | plastid | -0.8 *** | -0.5 ** | -0.5 * | 0.2 |
| *LHCB4.2* | AT3G08940 | g11705 | Light harvesting complex photosystem II | GO:0009536 | plastid | -0.8 * | -1.1 *** | -1.6 *** | -0.6 |
| *TAPX* | AT1G77490 | g01307 | Thylakoid ascorbate peroxidase | GO:0009536 | plastid | -0.7 ** | -0.8 *** | -0.9 * | -0.2 |
| *PSRP6* | AT5G17870 | g13318 | Plastid-specific 50S ribosomal protein 6 | GO:0009536 | plastid | -0.6 ** | -0.7 *** | -0.6 * | 0.0 |

[1]GO terms shown in Fig. 3a, b.

**Table S11.** Cd responsive genes in both populations from NM sites, which are also constitutively differentially expressed in Noss compared to either Pais or Wall.

| | | | | | | Log₂FC | | | |
|---|---|---|---|---|---|---|---|---|---|
| **Expression** | **Tissue** | **Name** | **AGI** | **Aha Id.** | **Description** | **Noss vs. Pais (-Cd)** | **Noss vs. Wall (-Cd)** | **+Cd vs. -Cd in Pais** | **+Cd vs. -Cd in Wall** |
| Noss < Pais/Wall | Root | NA | AT4G22640 | g14411 | Bifunctional inhibitor/lipid-transfer protein/seed storage 2S albumin superfamily protein | -0.8 * | -0.8 ** | -1.0 ** | -0.7 * |
| Noss < Wall | Root | *ACO2* | AT4G26970 | g04567 | aconitase 2 | 0.1 | -0.5 * | -0.8 ** | -1.0 *** |
| Noss < Pais | Shoot | *RTP1* | AT1G70260 | g03763 | Resistance to *Phytophthora parasitica* 1 | -1.1 * | 0.5 | -1.4 ** | -1.4 ** |
| Noss < Pais/Wall | Shoot | NA | AT5G02540 | g19446 | NAD(P)-binding Rossmann-fold superfamily protein | -2.2 *** | -2.0 *** | -1.7 * | -0.9 * |
| Noss < Wall | Shoot | NA | AT3G61820 | g05417 | Eukaryotic aspartyl protease family protein | 0.0 | -0.7 * | -1.5 *** | -1.2 *** |

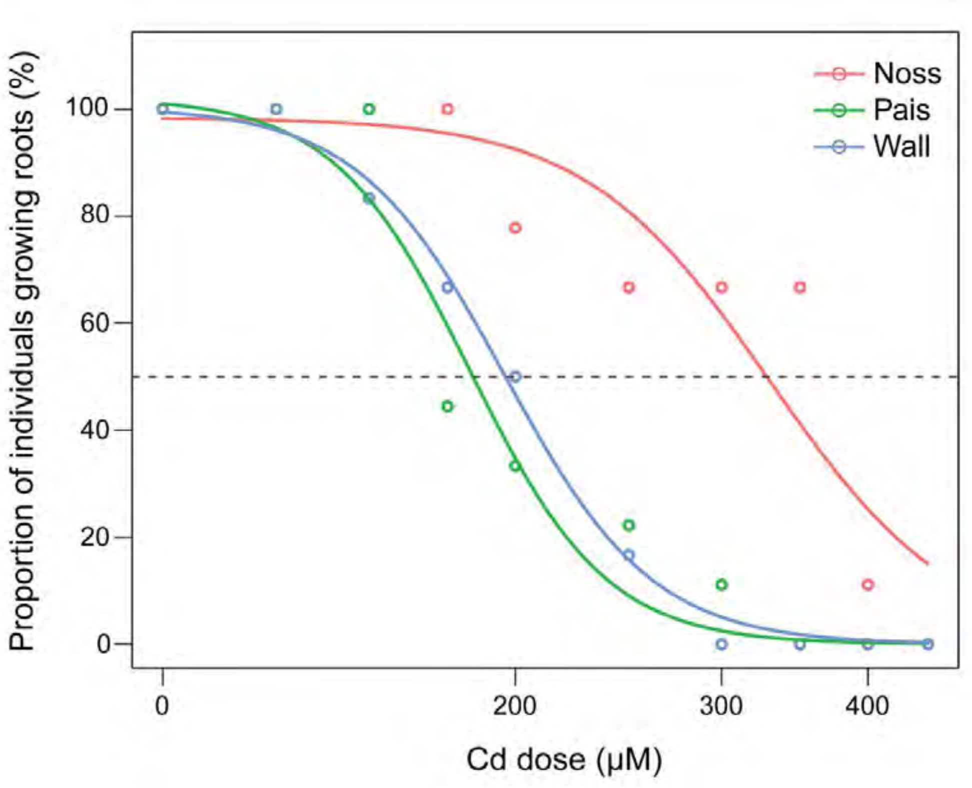

**Fig. S1.** Dose-response curve for the effect of Cd on root growth in the three *A. halleri* populations. Shown is the proportion of individuals maintaining root elongation as a function of Cd dose fitted to the tree-parameter generalized log-logistic model (see Fig. 1, Table S2). Datapoints (circles) represent the measured values used for model (lines) fitting. The dashed horizontal line indicates the proportion of 50% corresponding to $ED_{50}$ in the dose-response model.

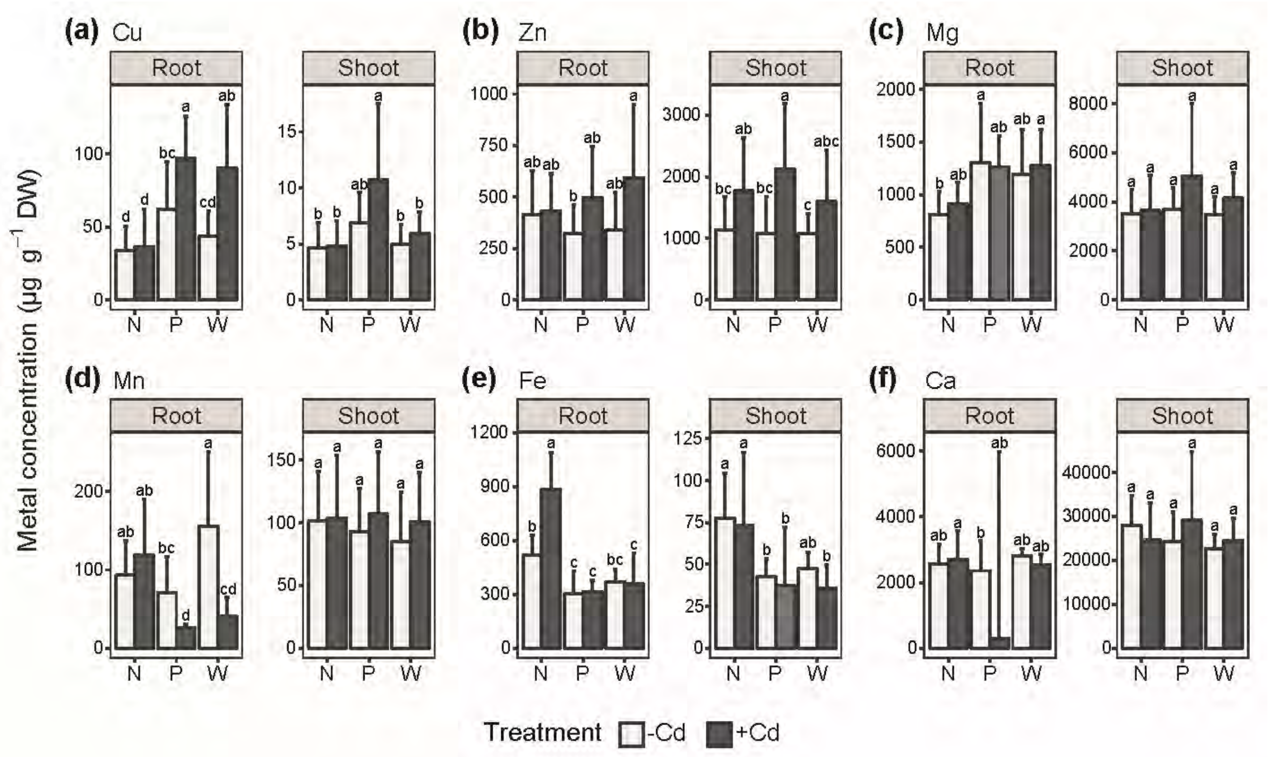

**Fig. S2.** Tissue metal concentrations in Cd-exposed and unexposed plants. (a-f) Concentrations of Cu (a), Zn (b), Mg (c), Mn (d), Fe (e) and Ca (f) in root and shoot tissues of hydroponically cultivated *A. halleri* originating from the Noss (N), Pais (P) and Wall (W) sites. Shown are mean ± SD ($n$ = 12 to 20 clones per population comprising both genotypes, from all three independent experiments). Four-week-old vegetative clones were exposed to 0 (-Cd) and 2 µM $CdSO_4$ (+Cd) in hydroponic culture for 16 d alongside the plants cultivated for transcriptome sequencing. Different characters denote statistically significant differences between means of log-transformed data based on two-way ANOVA, followed by Tukey's HSD test ($P < 0.05$). DW: Dry biomass.

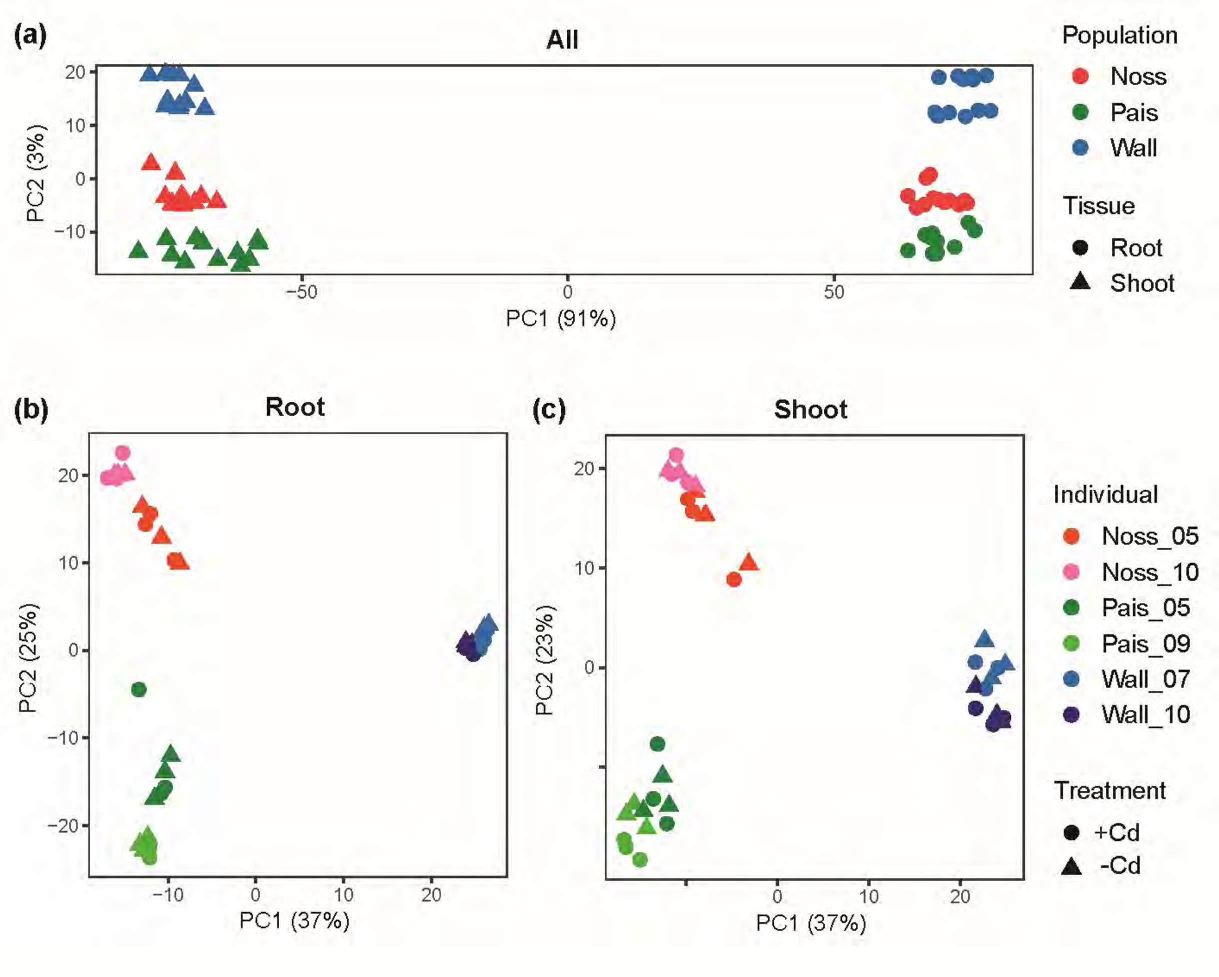

**Fig. S3.** Principal component analysis (PCA) of transcriptome data. (a-c) PCA of all data (*n* = 72, (A)), root (*n* = 36, (b)) and shoot (*n* = 36, (c)) (see Table S4). Variance stabilizing transformations of count values were used for PCA analysis. The percentages of variation attributable to PC1 and PC2 are given in parentheses.

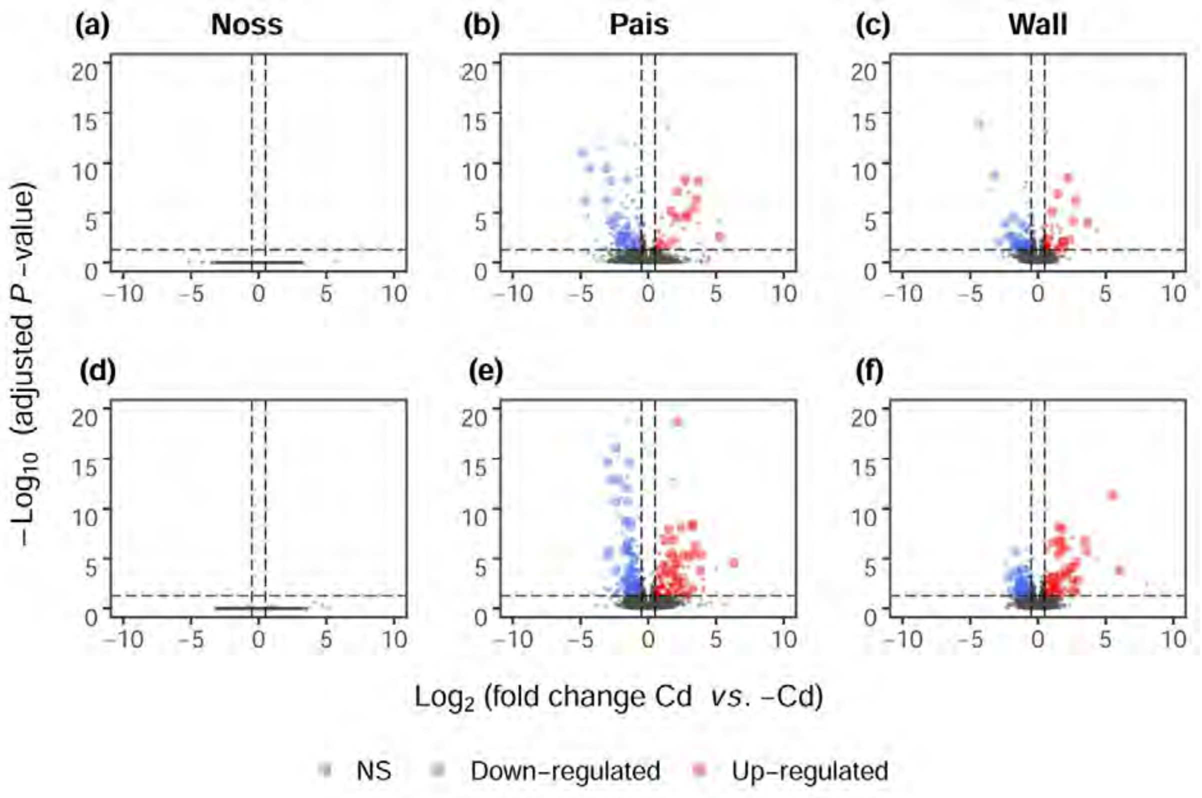

**Fig. S4.** Volcano plots of transcriptional responses to Cd in each population. (a-f) Data for root (a-c) and shoot (d-f) tissues of the Noss (a, d), Pais (b, e) and Wall (c, f) populations. Each datapoint represents the mean of three independent experiments, each conducted with two genotypes per population, for one gene. Up-regulated, $Log_2$(fold change +Cd *vs*. Control) > 0.5 and adjusted *P*-value < 0.05; down-regulated, $Log_2$(fold change +Cd *vs*. Control) < -0.5 and adjusted *P*-value < 0.05 (blue); NS, no significant change. Datapoints corresponding to entries of Table S10 are shown as larger and lighter colored symbols. Two datapoints, g27931 (*FER1*; $Log_2$(fold change) = -1.93, adjusted *P*-value = 6.80e-40) in (b) and g00781 (*CYP82C4*; $Log_2$(fold change) = -6.15, adjusted *P*-value = 3.76e-39) in (c) are not shown here.

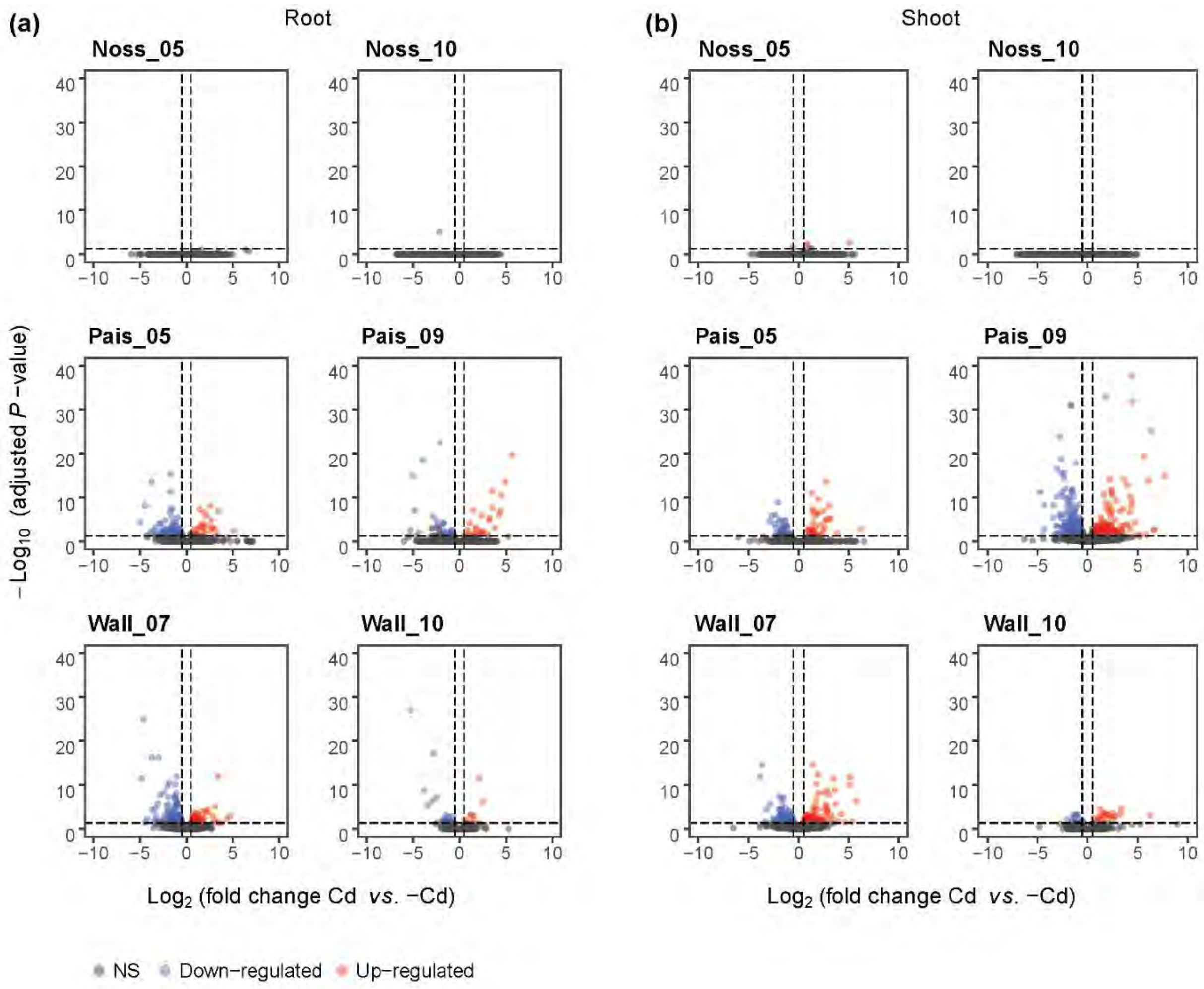

**Fig. S5.** Volcano plots of transcriptional responses to Cd in each genotype. (a, b) Data for root (a) and shoot (b) tissues shown for each of the genotypes. Each datapoint represents the mean of three independent experiments for one gene. Up-regulated, $Log_2$(fold change +Cd *vs*. Control) > 0.5 and adjusted *P*-value < 0.05; down-regulated, $Log_2$(fold change +Cd *vs*. Control) < -0.5 and adjusted *P*-value < 0.05; NS, not significant. Datapoint g00781 (*CYP82C4*; $Log_2$(fold change) = -7.39, adjusted *P*-value = 1.17e-64) for Wall_07 was omitted in (a).

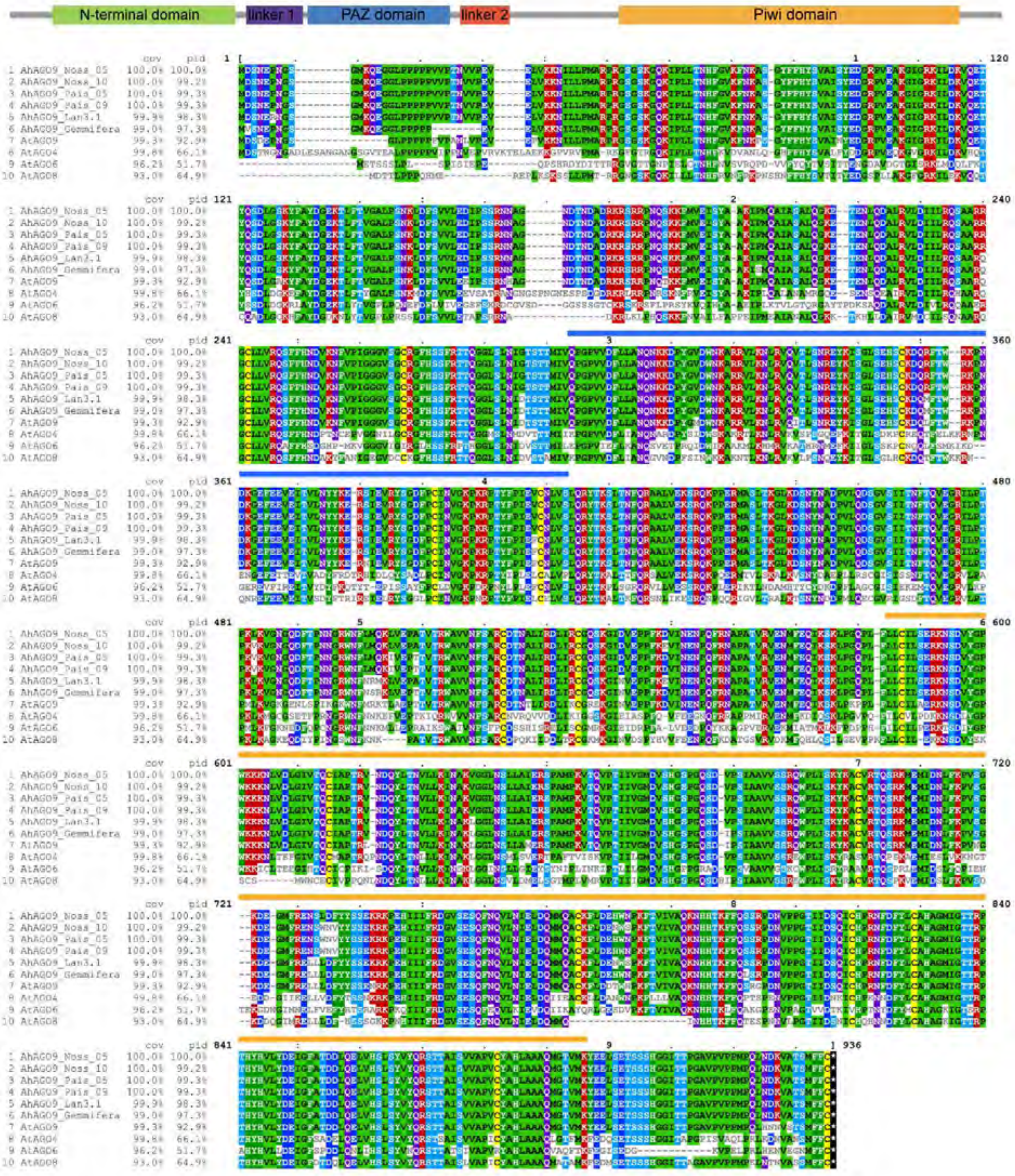

**Fig. S6.** Amino acid sequence alignment of AGO proteins. Shown are AGO9 amino acid sequences of *A. halleri* (Ah) individuals from the populations Noss and Pais, of which transcriptome sequencing was carried out in this study, and of *A. thaliana* (At) clade 3 ARGONAUTE family proteins AGO4, -6, -8 and -9 (cov, coverage; pid, percent identity). Positions of domains are shown by coloured bars: blue for PAZ (Piwi, Argonaute, and Zwille/pinhead) domain, yellow for Piwi (P-element Induced WImpy testis) domain.

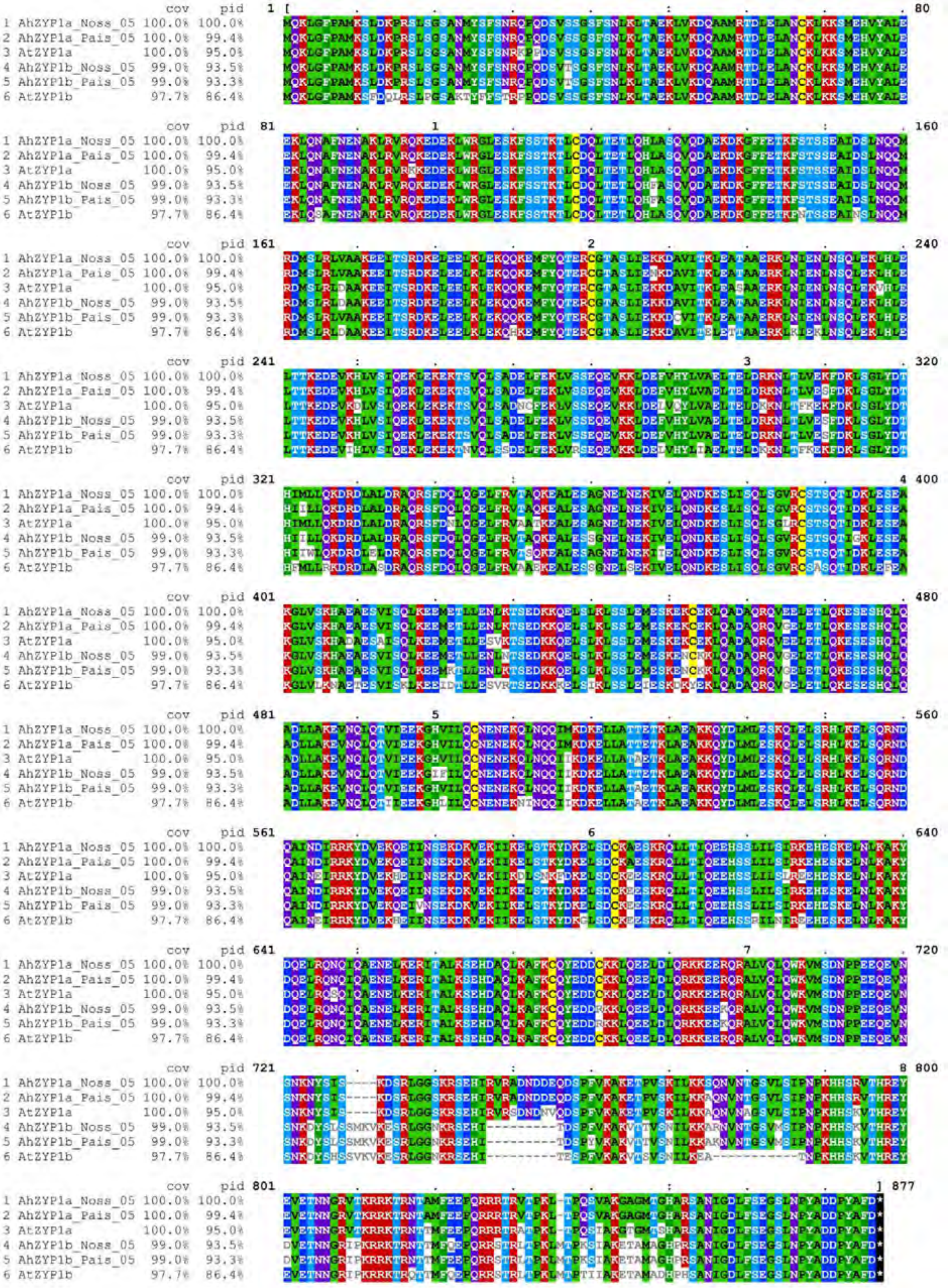

**Fig. S7.** Amino acid sequence alignment of ZYP1a/b proteins. Shown are amino acid sequences of *A. halleri* (Ah) individuals from the populations Noss and Pais, of which transcriptome sequencing was carried out in this study, and of *A. thaliana* (At) ZYP1a/b (cov, coverage; pid, percent identity).

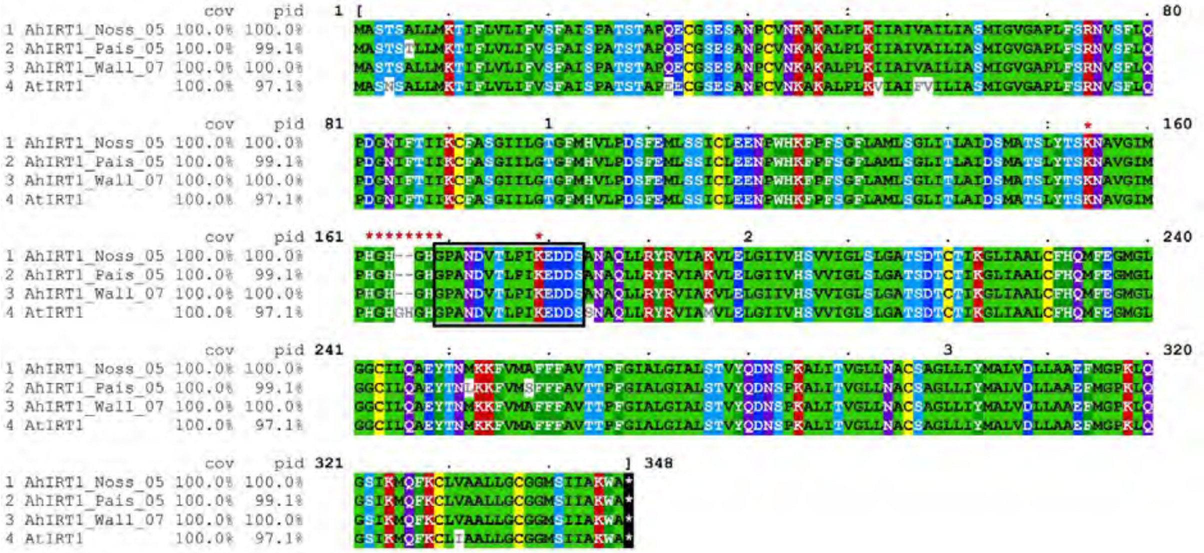

**Fig. S8.** Amino acid sequence alignment of IRT1 proteins. Shown are IRT1 amino acid sequences of *A. halleri* (Ah) individuals from Noss_05, Pais_05 and Wall_07, of which transcriptome sequencing was carried out in this study, and of *A. thaliana* (At) IRT1 (cov, coverage; pid, percent identity). Sequences below asterisks indicate a histidine (His) motif (HGHGHG) that potentially binds metals and two lysine (K) residues which might serve as attachment sites for ubiquitin. Sequences in the black box are the region corresponding to a synthetic peptide employed for the generation of the anti-*At*IRT1 ployclonal antibody in rabbits.

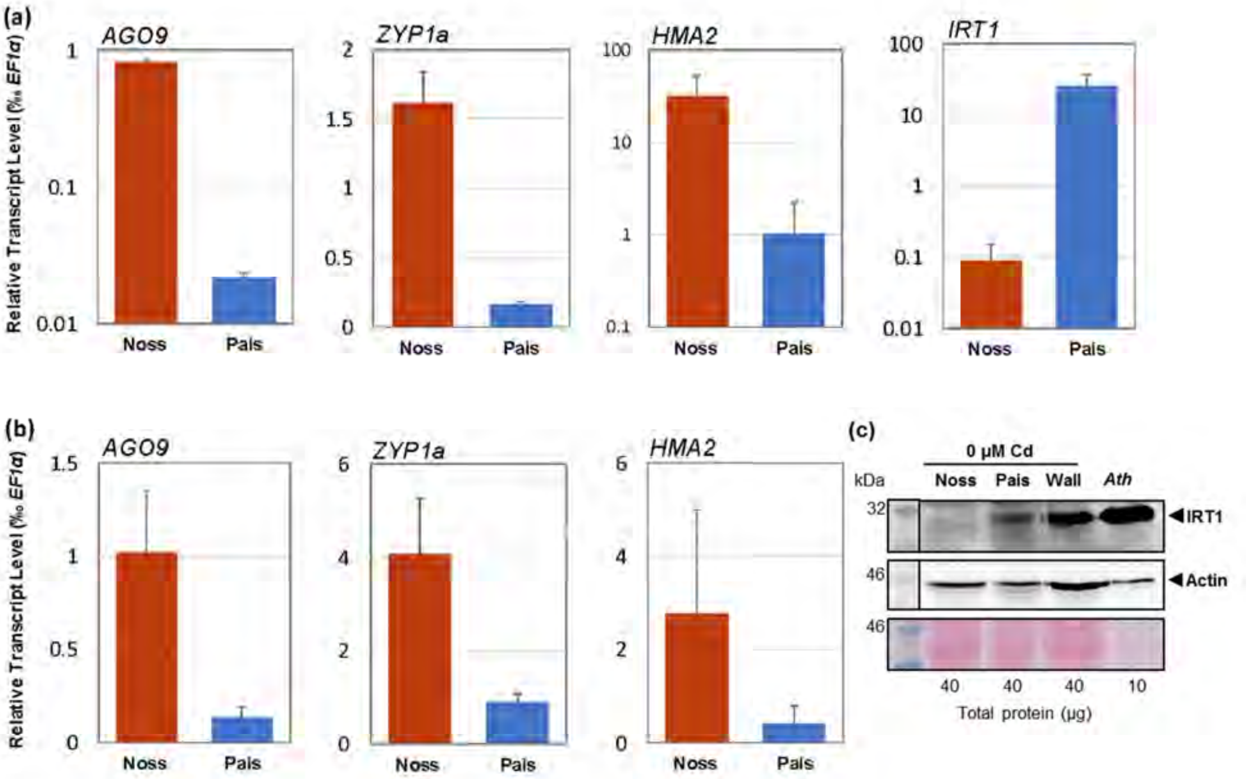

**Fig. S9.** Validation of sequencing-based transcriptomics (see Fig. 3). (a, b) Relative transcript levels of chosen candidate genes in roots (a) and shoots (b) of Noss and Pais as quantified by RT-qPCR. (c) Immunoblot of IRT1 (independent repeat). Total protein extracts (40 µg) from roots of Noss_05, Pais_09 and Wall_07 were separated on a denaturing polyacrylamide gel, blotted, and detection was carried out using an anti-IRT1 (top) or a n anti-Actin (center) antibody. The Ponceau S-stained membrane (after blotting) is shown at the bottom. Total protein extract (10 µg) from roots of Fe- and Zn-deficient *A. thaliana* (Ath, right lane) served as a positive control, with IRT1 detected as a single band at *ca*. 31 kDa. Shown are mean ± SD (*n* = 4, two genotypes per population and two independently synthesized cDNAs from one RNA extraction per genotype and one experiment), with each PCR reaction conducted in triplicate (a, b).

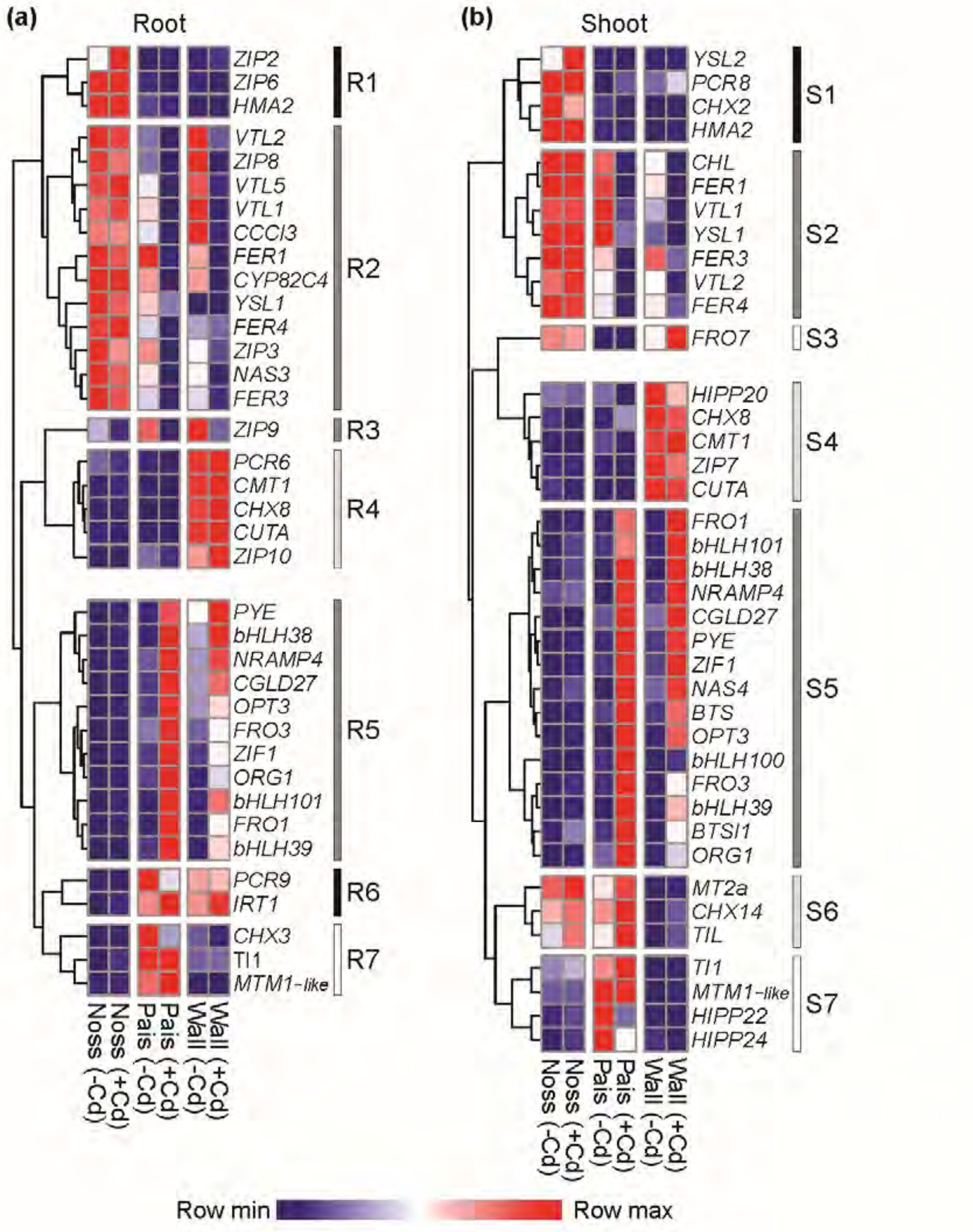

**Fig. S10.** Cluster analysis of metal homeostasis genes expressed differentially between populations and Cd treatments. (a, b) Heatmaps visualizing results from Euclidean distance clustering of relative transcript levels for roots (a) and shoot tissues (b). Shown are all metal homeostasis genes (Dataset S1) that were differentially expressed either between Cd and control treatments in any population, or between populations ($|Log_2$(fold change)$|$ > 1, adjusted *P*-value < 0.05, mean of normalized counts across all samples > 2; see Table S7). Normalized counts were averaged per population and treatment (*n* = 6). Subsequently, the row minimum was subtracted from each value. Finally, all values in each row were normalized to the row maximum. Clustering was conducted separately for root (a) and shoot (b) data. Clusters (R1-R7 and S1-S7) are marked by vertical lines (black, differentially expressed in Noss from both populations of NM soil origin (R1, R6 and S1); dark grey, Cd responsive genes in populations of NM origin (R2, R3, R5, S2 and S5); light grey, more highly (or less) expressed gene in Wall exclusively (R4, S4, S6); white, highly or less expressed in Pais exclusively (R7, S3, S7).

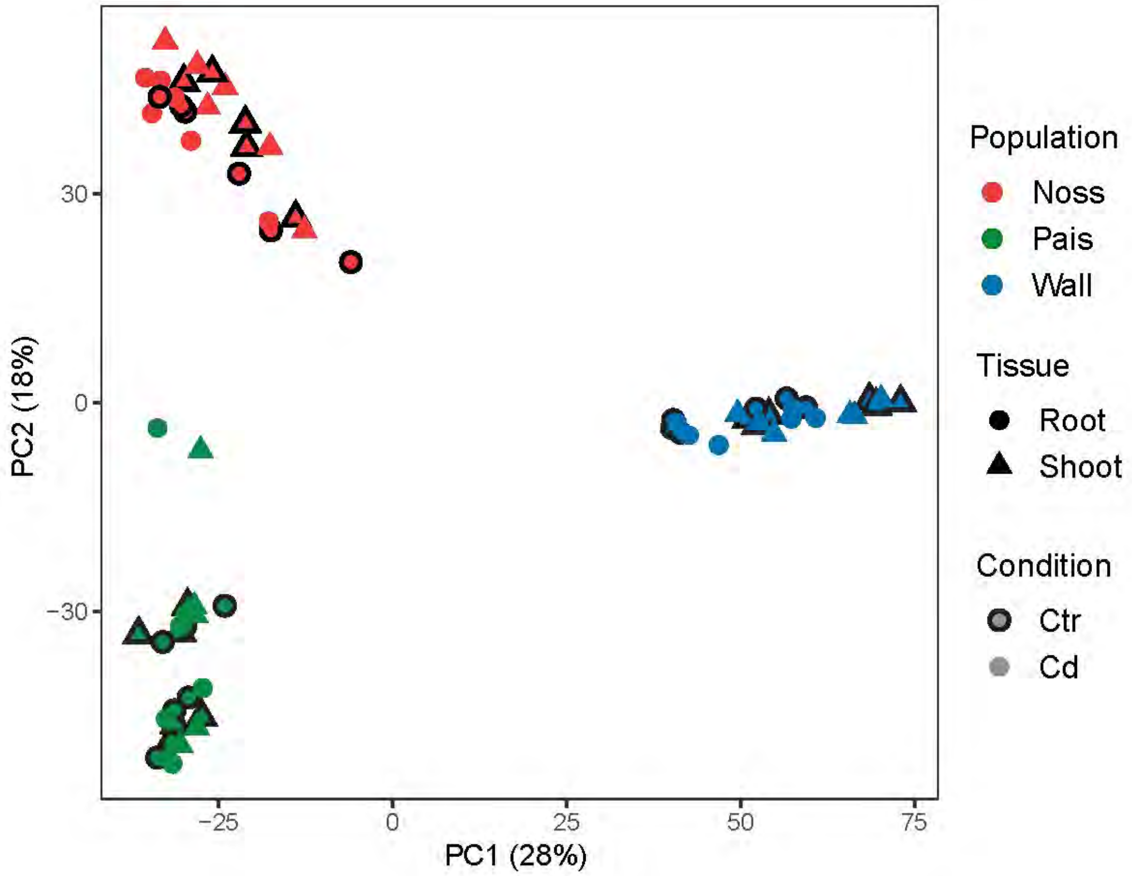

**Fig. S11.** PCA of transcripts derived from transposable element (TE) loci. Shown is a PCA of all samples (*n* = 72). Variance stabilizing transformations of normalized count values were used for PCA analysis. The percentages of variation attributable to PC1 and PC2 are given in parentheses.

### Supplemental Materials and Methods

#### Plant material and growth conditions

Plant individuals were taken as vegetative plants from three natural populations of *Arabidopsis halleri* ssp. *halleri* O'Kane & Al-Shehbaz, Noss and Pais in Italy, and the Wall population in Germany (specified in Table S1)(Stein *et al.*, 2017), as well as Laut (Germany) and Mias and Zapa (Poland) for a subset of experiments (Stein *et al.*, 2017), and were maintained in our glasshouse through successive generations of vegetatively propagated clones for at least 3 years. For all experiments, we produced vegetative cuttings of comparable sizes (six to nine leaves) from mother plants cultivated on standard soil (Einheitserde Minitray, Sinntal-Altengronau, Germany) in a growth chamber in 10-h light (90 µmol photons $m^{-2}$ $s^{-1}$, 22°C)/ 14-h dark (18°C) cycles at 65% constant relative humidity. Cuttings were rooted and pre-cultivated hydroponically in 0.2x modified Hoagland's solution (see below) under the same cultivation conditions for 4 weeks, except for the RT-qPCR analysis of AGO9 transcript levels in diverse accessions, we rooted vegetative stolons in 1:1 (v/v) peat:sand mix (round pots, 5 cm Ø, 3.5 cm depth, 50 mL volume) in a growth chamber for 17 days (20°C/17°C, 10 h light at 100 µmol $m^{-2}$ $s^{-1}$; GroBank BB-XXL.3, CLF Plant Climatics, Wertingen, Germany).
Subsequently, clones were transferred into 1x modified Hoagland's solution (1.5 mM $Ca(NO_3)_2$, 0.14 mM $KH_2PO_4$, 0.75 mM $MgSO_4$, 1.25 mM $KNO_3$, 25 µM $H_3BO_3$, 5 µM Fe-HBED, 5 µM $ZnSO_4$, 5 µM $MnSO_4$, 0.5 µM $CuSO_4$, 0.1 µM $Na_2MoO_4$, 50 µM KCl, 3 mM MES-KOH, pH 5.7)(Hoagland and Arnon, 1950, Becher *et al.*, 2004) and cultivated with weekly exchanges of solutions. For Cd tolerance assays, clones were sequentially exposed to stepwise increasing concentrations of $CdSO_4$ once per week (see Materials and Methods). Twenty plants were cultivated per container of 0.39 x 0.29 x 0.12 m (height) in 11 L of hydroponic solution in a growth chamber under the same conditions as employed for the cultivation of vegetative clones. For transcriptome sequencing and multi-element analysis, hydroponic media were supplemented with 2 or 0 (control) µM $CdSO_4$, respectively (see Table S4) for 16 d. Three plants were cultivated per round vessel of 0.125 m diameter, 0.11 m height, in 1 L of hydroponic solution. For the RT-qPCR analysis of *AGO9* transcript levels in diverse accessions, three plants were cultivated per 400-mL vessels of 400 ml hydroponic solution as above and supplemented with 10 nM $Pb(NO_3)_2$ and 0.25 µM $CdCl_2$, with two replicate vessels per genotype, for 3 w. During the final experimental phase until harvest, all plants were cultivated in 16-h light (90 µmol photons $m^{-2}$ $s^{-1}$, 22°C)/ 8-h dark (18°C) cycles at 60% constant relative humidity.

#### Cd tolerance assay

Root length of each clone cultivated under sequentially increasing Cd exposure was measured weekly and Cd tolerance index was quantified (see Materials and Methods). The proportions of individuals sustaining root elongation growth when exposed to stepwise increasing $CdSO_4$ doses were fitted to a dose-response model using the *drc* package in R (Ritz *et al.*, 2015) to estimate the Cd tolerance for the three populations alongside the $EC_{100}$. The three-parameter generalized log-logistic model function with lower limit 0 was used as follows:

$$f\big(x,\ (b,d,e)\big) = \frac{d}{1 + \exp\big(b(\log(x) - \log(e))\big)}$$

where $b$ denotes the slope parameter (coefficient denoting the steepness of the dose-response curve), $d$ represents the upper asymptotes or limits of the response and $e$ is the effective dose ($ED_{50}$) required to reach a proportion of 50% in a dose-response model.

**Harvest for multi-element analysis and transcriptome sequencing**

To determine the mineral composition of root and shoot tissues and for transcriptome sequencing, 6 to 10 and six replicate vegetative clones, respectively, were harvested per genotype condition and experiment at ZT7 (7 h after lights on). For multi-element analysis, freshly harvested roots of each plant were incubated in 30 mL ice-cold desorption solution I (2 mM $CaSO_4$, 10 mM $Na_2EDTA$, 1 mM MES-KOH pH 5.7) for 10 min, and then in 30 mL desorption solution II (0.3 mM bathophenanthroline disulphonate, 5.7 mM sodium dithionite) for 3 min (Cailliatte *et al.*, 2010). Roots were then rinsed in 100 ml ice-cold $ddH_2O$ with gentle stirring for 1 min. Subsequently, both roots and shoots were washed twice in 100 mL $ddH_2O$. After blotting dry with tissue paper, the samples were placed into paper bags and left to dry in ambient air at room temperature. For transcriptome sequencing, roots and shoots were separated using a razor blade, roots were briefly blotted dry on tissue paper, and root and shoots were placed in pre-chilled 50-mL screw-cap polypropylene tubes, frozen in liquid nitrogen and stored at -80°C.

**Multi-element analysis**

Tissues were dried additionally at 60°C for 3 to 5 d and subsequently equilibrated in ambient air for ≥ 24 h, then ground to a fine powder using ceramic beads in a Precellys 24 at 6,500 rpm for 20 s (Bertin Technologies, Montigny le Bretonneux, France). Subsamples of *ca.* 20 mg dried tissue powder were used for microwave digestion, followed by multi-element analysis using Inductively-Coupled Plasma Optical Emission Spectrometry (ICP-OES; iCAP 6500 Duo; Thermo Scientific, Waltham, USA), both conducted as described previously (Stein *et al.*, 2017).

**Preparation of RNA libraries and sequencing**

In each experiment, root and shoot tissues were pooled separately from all six clones cultivated per genotype and treatment, and homogenized in liquid nitrogen using a pestle and mortar. Total RNA was extracted from aliquots of 50 to 100 mg per sample using the RNeasy Plant Mini Kit (Qiagen, Hilden, Germany) according to the manufacturer's instructions. RNA was quantified using an EON Microplate Spectrophotometer (BioTek, Bad Friedrichshall, Germany). For library preparation, 1 µg of total RNA per sample was used utilizing NEBNext Ultra II RNA Library Prep Kit for Illumina and NEBNext Multiplex Oligos for Illumina (Dual Index Primer Set 1) according to the manufacturer's instructions (New England Biolabs, Frankfurt, Germany). Quality and quantity (≥ 0.5 ng $\mu l^{-1}$) of libraries were verified using an Agilent 2100 Bioanalyzer (Agilent, Santa Clara, CA, USA) and a Qubit 2.0 Fluorometer (Thermo Fisher Scientific, Schwerte, Germany). Sequencing was conducted on the Illumina® HiSeq X Ten platform; after removing all reads containing adapters, > 10% N (i.e., base identity not determined) or Phred Q score < 10 in > 50% of bases, we obtained 14 to 29 mio. 150-bp read pairs per sample (Novogene, Hongkong, China). All read data are available here (PRJEB35573, ENA, EMBL-EBI).

**Transcriptome sequencing data analysis**

Adequate average per base read quality, duplication level, GC content and length distribution were ensured using FastQC (http://www.bioinformatics.babraham.ac.uk/projects/fastqc/). Subsequently, all reads were aligned to the reference genome sequence of *A. halleri* ssp. *gemmifera* (Matsumura) O'Kane & Al-Shehbaz accession Tada mine (W302) (Briskine *et al.*, 2017) using HISAT2 version 2.1.0 (Kim *et al.*, 2015), with options set to default except not writing unaligned reads to the output (--no-unalign). Multiply mapping reads were then processed using the error correction scripts according to step 3 of COMEX 2.1 (Pietzenuk *et al.*, 2016) for those reads aligning multiple times with either the same start or the same end position, or palindromic reads, i.e. a single read mapping on opposing DNA strands in an identical position. To avoid read count bias, only the shortest alignments were retained (same start or same end) and one alignment at random for each palindromic multiple mapping read. Thereafter, the number of fragments mapped per gene were determined using Qualimap2 (comp-counts) with the options, -p non-strand-specific; -algorithm proportional (Okonechnikov *et al.*, 2016). Fragment counts per gene for each sample were used for principal component analysis (PCA) and differential gene expression analysis using the R package *DESeq2* (Love *et al.*, 2014). Differentially expressed genes (DEGs) between the Cd treatment and the control condition (2 *vs*. 0 µM $CdSO_4$) were identified upon per-tissue per-population grouping of count data for normalization. For the identification of DEGs between populations, normalization was done across all samples per tissue, because library sizes and the distribution of normalization factors differed between root and shoot samples. All *P*-values (Wald-Test) were adjusted for FDR correction (Benjamini and Hochberg, 1995). In addition to the standardized outputs, normalized counts extracted from the results of *DESeq2* were divided by each gene length (NCPK; normalized counts per kilobase of gene length) for the representation and comparison of transcript abundance across both samples and genes. The significances for the probabilities of the intersections in the Venn diagrams for the numbers of DEGs were calculated using the hypergeometric distribution through the R package *gmp* (R_Core_Team, 2013). Gene clustering based on Euclidean distance was performed using the R package *pheatmap* after selection of DEGs between conditions and all population pairs based on the genes contained in our manually curated custom annotated list of metal homeostasis-related genes (Dataset S1, see column K). For gene ontology (GO) term enrichment analyses *A. halleri* gene IDs were converted into *A. thaliana* TAIR10 AGI codes (Briskine *et al.*, 2017), which were then used in hypergeometric distribution estimation in the function of g:GOSt built in g:Profiler (Reimand *et al.*, 2016).

**Quantitative RT-qPCR validation of transcript levels as determined by RNA-seq**

Total RNA was extracted from aliquots of the same tissue homogenates as used for transcriptome sequencing (-Cd controls) using the RNeasy Plant Mini with DNase I treatment during extraction according to the manufacturer's instructions (Qiagen). RNA quality was verified spectrophotometrically (NanoDrop 2000 Spectrophotometer, Thermo Fisher Scientific), and its integrity was verified in a non-denaturing 1% (w/v) agarose gel. One µg of DNase-treated RNA was used in a 20-µL first-strand cDNA synthesis reaction using RevertAid First Strand cDNA Synthesis Kit (Thermo Fisher Scientific) containing 1 µl of RevertAid M-MuLV RT enzyme, 1 µl of RiboLock RNase Inhibitor, 2 µl of 10 mM dNTP Mix and 1 µl of oligo (dT) primer, with incubation at 42°C for 60 min according to the manufacturer's instructions. Real-time RT-PCR reactions were performed in a 384-well LightCycler®480 II System (Roche Diagnostics, Mannheim, Germany). Equal amounts of cDNA corresponding to approximately 10 ng of mRNA were used in each reaction containing 4 µL of a 1/20 dilution of the synthesized cDNA, 5 µL GoTaq qPCR Mastermix (Promega, Walldorf, Germany) and 1 µL of 2.5 µM forward and 2.5 µM reverse primer mix in a total volume of 10 µL. The thermal profile was: initial denaturation at 95°C for 5 min, 40 cycles at 95°C for 1 min and 60°C for 1 min, followed by final elongation at 60°C for 10 min, followed by melting curve analysis starting at 95°C with a ramp rate of -0.11°C $s^{-1}$ in order to verify the amplification of a single PCR product per well. For the quantification of relative transcript levels (RTL), the reaction efficiencies (RE) and cycle thresholds ($C_T$, mean of six technical replicates) were determined using LinRegPCR (Ramakers *et al.*, 2003). Reactions with primer amplification efficiencies below 1.8 were discarded, and mean primer amplification efficiency (PE) for each primer pair calculated from all reactions. Transcript levels of genes of interest within a cDNA were normalized to the respective transcript level of *EF1α* or *Helicase* (*HEL*) as a constitutively expressed control gene (*CCC*) using the following formula: $\mathbf{RTL} = (\mathbf{PE}_{CCC}{}^{C_{T\,CCC}}) \cdot (\mathbf{PE}_{target}{}^{-C_{T\,target}})$(Talke *et al.*, 2006)(See Table S8 for primer sequences).

**Immunoblots**

For immunodetection of IRT1, 100-mg aliquots of powdered frozen root tissue (as harvested for RNA-seq) were extracted in 300 µL of extraction buffer (50 mM TRIS-HCl pH 8.0, 25 mM EDTA, 5% (w/v) SDS, 2% (w/v) DTT, 0.1% (w/v) orthophenantroline and 1 mM phenylmethylsulfonyl fluoride) (Seguela *et al.*, 2008). After vortexing at maximum speed for 20 s, samples were incubated for 5 min at room temperature to solubilize membranes and cell debris were pelleted by centrifugation at 14,000 g, 4°C for 15 min. Protein concentration in the supernatant (1:5 dilution) was determined with the Pierce™ BCA Protein Assay Kit (Thermo, USA) using BSA as a standard (25 to 2,000 µg $ml^{-1}$) and 1:5 diluted buffer as a blank. Forty µg of total protein (10 µg for the positive control, roots of *A. thaliana* grown as described (Talke *et al.*, 2006) with omission of FeHBED from the hydroponic solution during the final week of cultivation before harvest) was separated on a 12.5% (v/v) SDS-PAGE gel (Lämmli, 1970) at 20 mA and room temperature (RT), followed by wet/tank transfer to nitrocellulose membranes (4°C, 100 V, 1 h). Blots were blocked in TRIS-Buffered Saline (20 mM TRIS Base, 150 mM NaCl, pH 7.6) with 0.05% (v/v) Tween-20 (TBST) and 5% (w/v) non-fat milk at RT with horizontal agitation at 60 rpm for 1 h. The membrane was incubated with the primary antibody (anti-IRT1, AS11 1780, Lot number 1203; Agrisera, Vännas, Sweden) diluted 1:500 in TBST with 2.5% (w/v) non-fat milk (TBSTM) at RT for 1 h. The membrane was rinsed briefly 3 times, washed in TBST for 10 min, then incubated with an HRP-conjugated secondary antibody (31466; Lot number RL243150; Thermo Fisher Scientific) diluted 1:10,000 in TBSTM. After three brief rinses and a wash in TBST for 10 min, signal development was carried out at RT for 5 min with ECL Select Western Blotting Detection Reagent (GE Healthcare, Little Chalfont, England). The blot was imaged with a Fusion Fx7 GelDoc (Vilber Lourmat, Eberhardzell, Germany) with 30 s exposure. Following detection, the PVDF membrane was stained with Coomassie Blue (0.025% (w/v) Coomassie brilliant blue R-250 in 50% methanol/10% acetic acid (v/v)) for 5 min and de-stained with 40% methanol/10% acetic acid (v/v) for 5-10 min. Coomassie-stained membrane served as control for sample loading. For ACTIN detection, membranes were re-probed after mild stripping. For stripping, membranes were incubated at RT twice in stripping buffer (1.5% (w/v) glycine, 0.1% (w/v) SDS, 1% (v/v) Tween-20) for 10 min and subsequently washed twice in Phosphate-Buffered Saline (PBS; 137 mM NaCl, 2.7 mM KCl, 8 mM $Na_2HPO_4$, and 2 mM $KH_2PO_4$) (10 min) and twice in TBS with 0.1% (v/v) Tween-20 (TBST) (5 min). Blots were blocked in TBST with 5% (w/v) non-fat milk as described above. The membrane was probed with an anti-ACTIN antibody (AS13 2640, Lot Number 1203; Agrisera) diluted 1:2,500 in TBSTM (2% (w/v) non-fat milk) at RT for 1 h and with Goat anti-Rabbit IgG (H+L) HRP-conjugated secondary antibody diluted 1:10,000 in TBSTM (31466; Lot number RL243150; Thermo Fisher Scientific) at RT for 1 h.

**Analysis of transposable element (TE) transcript levels**

To analyze the transcriptional activity of TEs in the three populations, the Illumina RNA-seq reads were aligned to the *Arabidopsis lyrata* reference genome assembly (Hu *et al.*, 2011) using HISAT2 version 2.1.0 (Kim *et al.*, 2015) because of the substantially higher quality of this assembly outside protein-coding regions. Mapped reads were then corrected using the complete COMEX 2.1 pipeline (Pietzenuk *et al.*, 2016) in two steps. First, multiple mapping error correction was performed, in which reads that mapped multiple times with same start or same end and palindromic (same position on opposing DNA strands) were handled as described above. Second, multiple mapping reads across different TE families were excluded from further analysis (Pietzenuk *et al.*, 2016). Subsequently, the numbers of reads per TE locus were retrieved from the output file of the COMEX2.1 using Qualimap2, with the same options as used in transcriptome sequencing data analysis (see above). Transposable element information used in COMEX 2.1 pipeline and read counting was based on the TE annotation of *A. lyrata* MN47 (Pietzenuk *et al.*, 2016). Transcript levels for each TE locus were approximated based on RPKM (Reads Per Kilobase per Million reads)(Mortazavi *et al.*, 2008). The global minimum threshold for considering a TE transcriptionally active was determined based on density plots (RPKM $\geq$ 7 per locus). Total transcript levels for each TE type were calculated by summation based on RPKM.

**Variants of *AGO9*, *ZYP1a/b* and *IRT1* cDNA sequences in *A. halleri***

We established the cDNA sequences of *AGO9*, *ZYP1a/b* and *IRT1* as one consensus sequence per genotype from all RNA-seq and gDNA reads (see also Table S6) that mapped to the corresponding locus in the *A. halleri* ssp. *gemmifera* reference genome assembly (Briskine *et al.*, 2017), employing the Integrated Genome Viewer (Robinson *et al.*, 2011, Thorvaldsdottir *et al.*, 2013, Robinson *et al.*, 2017) and Clustal Omega (Madeira *et al.*, 2019). Completeness and orthology of the constructed consensus *AGO9*, *ZYP1a/b* and *IRT1* coding sequence (cds) variants from different *A. halleri* genotypes were validated against *A. thaliana* Col-0 genomic and cds from The Arabidopsis Information Resource (TAIR; www.arabidopsis.org)(Berardini *et al.*, 2015). Following conceptual translation using ExPASy (https://web.expasy.org/translate/)(Gasteiger *et al.*, 2003), amino acid sequence alignments were generated using Clustal Omega and MView (Madeira *et al.*, 2019) of the variants identified here and orthologues in *A. thaliana* and *A. halleri* (*Arabidopsis halleri* v1.1, DOE-

JGI, http://phytozome.jgi.doe.gov/) (Figs. S6-S8). Subsequently, separate multiple cDNA sequence alignments were generated of *AGO9*, *ZYP1a/b* and *IRT1* from *A. halleri* populations for the visualization and identification of conserved regions, which were then used to design RT-qPCR primers using Primer3 (Koressaar *et al.*, 2018), followed by validation through the absence of predicted secondary structures using mfold (Zuker, 2003)(Table S8). All cds were deposited under repository number MN747968-78 (submission ID: 2287280, Genbank, NCBI).

### Other statistical data analysis

Multiple comparisons of means were conducted using two-way ANOVA and Nested ANOVA including an error term for genotype, followed by Tukey's HSD test, with the *stats* and *agricolae* package in R (R_Core_Team, 2013). For each statistical test, either original numbers, Log-transformed or Box-Cox-transformed numbers were used, depending on Shapiro-Wilk normality testing utilizing the *stats* package, and also based on homoscedasticity (equality of variances) by Levene's test using the *car* package in R. Transformation types were chosen for which *P*-value were closest to 1 in Shapiro-Wilk and Levene's tests.
